# Systematic benchmarking of imaging spatial transcriptomics platforms in FFPE tissues

**DOI:** 10.1101/2023.12.07.570603

**Authors:** Huan Wang, Ruixu Huang, Jack Nelson, Ce Gao, Miles Tran, Anna Yeaton, Kristen Felt, Kathleen L. Pfaff, Teri Bowman, Scott J. Rodig, Kevin Wei, Brittany A. Goods, Samouil L. Farhi

**Affiliations:** Spatial Technology Platform, Broad Institute of MIT and Harvard, Cambridge, MA 02142 USA; Thayer School of Engineering, Molecular and Systems Biology, and Program in Quantitative Biomedical Sciences at Dartmouth College, Hanover, NH 03755, USA; Division of Rheumatology, Inflammation, and Immunity, Brigham and Women’s Hospital and Harvard Medical School, Boston, MA 02215, USA; Immunai, New York, NY 10016, USA; ImmunoProfile, Brigham & Women’s Hospital and Dana-Farber Cancer Institute, Boston, MA 02215, USA; Center for Immuno-Oncology, Tissue Biomarker Laboratory, Dana-Farber Cancer Institute, Boston, MA 02215, USA; Department of Pathology, Brigham and Women’s Hospital, Boston, MA 02215, USA

## Abstract

Emerging imaging spatial transcriptomics (iST) platforms and coupled analytical methods can recover cell-to-cell interactions, groups of spatially covarying genes, and gene signatures associated with pathological features, and are thus particularly well-suited for applications in formalin fixed paraffin embedded (FFPE) tissues. Here, we benchmarked the performance of three commercial iST platforms on serial sections from tissue microarrays (TMAs) containing 23 tumor and normal tissue types for both relative technical and biological performance. On matched genes, we found that 10x Xenium shows higher transcript counts per gene without sacrificing specificity, but that all three platforms concord to orthogonal RNA-seq datasets and can perform spatially resolved cell typing, albeit with different false discovery rates, cell segmentation error frequencies, and with varying degrees of sub-clustering for downstream biological analyses. Taken together, our analyses provide a comprehensive benchmark to guide the choice of iST method as researchers design studies with precious samples in this rapidly evolving field.

## MAIN

Spatial transcriptomics (ST) tools measure the gene expression profiles of tissues or cells *in situ*. These approaches overcome the limitations of single-cell RNA-sequencing (scRNA-seq) methods by negating the need for cellularization and maintaining both local and global spatial relationships between cells within a tissue. ST can thus recover cell-cell interactions with high confidence, groups of spatially covarying genes, groups of cells predictive of cancer survival, and gene signatures associated with pathological features [1, 2]. These advantages, coupled with rapidly emerging computational and analytical methods, have led to substantial excitement about deploying these platforms in fundamental biology studies, and in the clinic for research and diagnostic purposes [3, 4, 5].

ST tools can be split into two broad categories: sequencing (sST) and imaging (iST) based modalities. sST methods tag transcripts with an oligonucleotide address indicating spatial location, most commonly by placing tissue slices on a barcoded substrate; isolating tagged mRNA for next-generation sequencing; and computationally mapping transcript identities to locations [6]. In contrast, iST methods most commonly use variations of fluorescence *in situ* hybridization (FISH) where mRNA molecules are tagged with hybridization probes which are detected in a combinatorial manner over multiple rounds of staining with fluorescent reporters, imaging, and de-staining **(Fig. 1a)** [7]. Computational reconstruction then yields maps of transcript identity with single-molecule resolution. Compared to sST methods, iST methods are targeted to subsets of the transcriptome due to their reliance on pre-defined gene panels and they adopt the higher spatial resolution and sensitivity of FISH, yielding single-cell resolution data [8].

**Figure 1:**
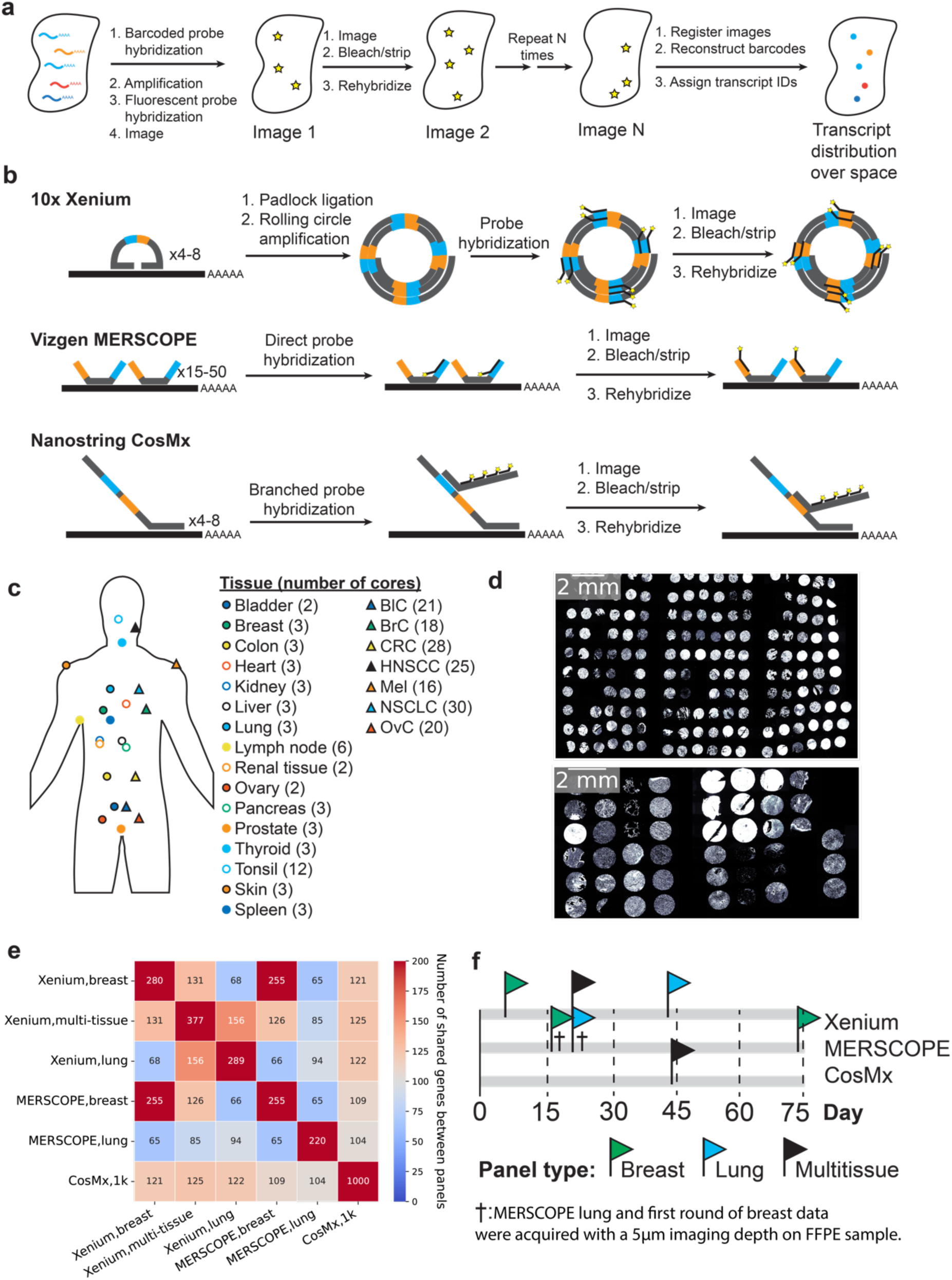
Experimental design and iST platforms. (a) Overall approach for generating iST data. (b) Different amplification approaches for Xenium, MERSCOPE, and CosMx. (c) Overview of the tissue types and numbers of cores used in this study. BlC = bladder cancer, BrC = breast cancer, CRC = colorectal cancer, HNSCC = head and neck squamous cell carcinoma, Mel = Melanoma, NSCLC = non-small cell lung cancer, OvC = ovarian cancer. (d) DAPI images from the Xenium run of each TMA, including tumors (top) and normal tissues (bottom) (e) The number of common target genes in each panel used in this study. (f) Overall timeline of the imaging days for each study. Day = 0 corresponds to the day of slicing. † denotes the MERSCOPE breast and lung panels acquired with a 5 µm imaging thickness, thinner than manufacturer instructions.

While the iST methods share some similarities, significant differences arise in primary signal detection and amplification, sample processing, and the subsequent fluorescent cycling chemistry (**Fig. 1b**) [9,10,11]. The need for amplification of signal is coupled to the sample processing, namely whether the sample is cleared, gel-embedded, or photobleached to quench autofluorescence. There are tradeoffs due to differences in sample processing for each iST method. For example, clearing of the sample increases signal quality but can prevent follow-up H&E staining and complicate immunostaining, which, in turn, can make cell segmentation more challenging. Finally, there are tradeoffs between imaging time, molecular plex, and imaging area covered, which result from the particular combination of the molecular protocol and the imaging hardware implementation [12].

A key historic limitation in the widespread use of iST methods with human clinical samples was the incompatibility of most methods with formalin-fixed, paraffin-embedded (FFPE) tissue samples [13, 14]. FFPE is the standard format for clinical sample preservation for pathology due to its ability to maintain tissue morphology and sample stability at room temperature for decades [15]. The ability to process FFPE samples with iST would enable the use of archival tissue banks for studies and obviate the need for specialized sample harvesting workflows. However, FFPE samples tend to suffer from decreased RNA integrity, particularly after having been stored in archives for extended periods of time [16].

Three companies recently released the first FFPE compatible commercial iST platforms: 10x’s Xenium, NanoString’s CosMx, and Vizgen’s MERSCOPE [9,10,11,17]. These three platforms each use different protocols, probe designs, signal amplification strategies, and computational processing methods, and therefore may potentially yield different sensitivities and downstream results. The main chemistry difference lies in transcript amplification: 10x Xenium uses a small number of padlock probes with rolling circle amplification; CosMx uses a low number of probes amplified with branch chain hybridization; and MERSCOPE uses direct probe hybridization but amplifies by tiling the transcript with many probes (**Fig. 1b**). However, no head-to-head performance comparisons on matched samples have been published. Understanding the key differences across platforms will allow users to make better-informed decisions regarding panel design, method choice, and sample selection as they design costly experiments, often on precious samples that have been bio-banked for years [18].

In this study, we compared currently available FFPE-compatible iST platforms on matched tissue samples. We prepared a set of samples representative of typical archival FFPE tissues, comprised of 23 different tissue types, and acquired matched data from sequential sections according to the manufacturer’s best practices at the time of writing, generating a dataset of > 3.3M cells. We analyzed the relative sensitivity and specificity of each method on shared transcripts, and further quantified the concordance of the iST data across each platform with orthogonal data sets from The Cancer Genome Atlas (TCGA) program and Genotype-Tissue Expression (GTEx) databases [19,20]. Then we focused on cell-level comparisons, evaluating the out-of-the-box segmentation for each platform based on detected genes and transcripts and coexpression patterns of known disjoint markers. Finally, we cross-compared the ability of each platform to identify cell type clusters with breast and breast cancer tissues as an example use case. Taken together, our work provides the first head-to-head comparison of these platforms across multiple archival healthy and cancerous FFPE tissue types.

## RESULTS

### Collection of matched iST data across 23 FFPE tissue types reveals high transcript counts obtained by Xenium and CosMx

To test the performance of the latest generation of FFPE-compatible iST tools, we measured the spatial expression of the same genes on the same samples as much as possible given current panel configurations. To accomplish this, we used two previously generated multi-tissue tissue microarrays (TMAs) from clinical discarded tissue (**see Methods**). We focused on FFPE tissues as the standard method for sample processing and archival in pathology. One TMA consisted of one hundred and seventy 0.6 mm diameter cores (i.e. sampled regions) from seven different cancer types, with 3-6 patients per cancer type, and 3-6 cores per patient (**Fig. 1c-d, Supplementary Table 1**). A separate TMA consisted of forty-five 1.2 mm diameter cores spanning sixteen normal tissue types isolated with each tissue type coming from one patient and represented in 2-3 cores (**Fig. 1c-d, Supplementary Table 2**). CosMx and Xenium suggest pre-screening samples based on H&E, while MERSCOPE recommends a DV_200_ > 60%. Since our goal was to determine the compatibility of iST platforms under typical workflows for biobanked FFPE tissues, and since TMAs are challenging to assay by DV_200_, samples were not prescreened based on RNA integrity. Samples were screened by H&E in the process of TMA assembly. Both TMAs were sliced into serial sections for processing by 10x Xenium, Vizgen MERSCOPE, and NanoString CosMx, following manufacturer instructions (**see Methods**).

The three different iST platforms offer different degrees of customizability and panel compositions. In terms of panel design, MERSCOPE and Xenium offer either fully customizable panels or standard panels with optional add-on genes, while CosMx offers a standard 1K (substantially larger plex) panel with optional add-on genes. We opted to run the CosMx 1K panel as available commercially, as well as the Xenium human breast, lung, and multi-tissue off-the-shelf panels. We then designed two MERSCOPE panels to match the pre-made Xenium breast and lung panels, by filtering out any genes which could potentially lead to high expression flags in any tissue in the Vizgen online portal. This resulted in a total of six panels, with each panel overlapping the others on > 94 genes (**Fig. 1e, Supplementary Table 3**). Samples were run following manufacturer instructions over the course of 74 days after slicing (**Fig. 1f, Supplementary Table 4**), with efforts made to ensure that head-to-head comparisons were available at similar time points for each pair of platforms. In data review, we noticed that MERSCOPE breast and lung panel were originally acquired with a 5 µm imaging depth, which was unintentionally thinner than the manufacturer recommendation of 10 µm, and could thus lead to aberrantly low counts. Thus, a second round of breast panel acquisition was performed with a 10 µm imaging depth (**Supplementary Table 1a**), resulting in a median 3.0-fold increase in expression across all transcripts. We excluded the 5 µm MERSCOPE breast panel data from all further comparisons but left the lung panel data in as an illustrative example of an unsuccessful run. However, we emphasize that MERSCOPE performance should be judged based on the rerun breast panel.

Each data set was processed according to the standard base-calling and segmentation pipeline provided by each manufacturer. The resulting count matrices and detected transcripts were then subsampled and aggregated to individual cores of the TMA (**Methods**). Across all datasets we generated >190 million transcripts, >3.3 million cells, across 7 tumor types, and 16 normal tissue types. Overall, we found that the cores from each TMA were generally well adhered to the tissue and detected transcripts, and we were able to collect data from all three modalities for 217 cores (**Supplementary Table 4**). The total number of transcripts recovered for each run was highest for Xenium, followed closely by CosMx, and then MERSCOPE (**Supplementary Table 4**). The total number of cells initially reported was highest for Xenium followed by MERSCOPE and CosMx (**Supplementary Table 4**). Based on the initially reported number of transcripts, the tumor TMA appeared to provide more counts than the normal tissue TMA, which we ascribed to a higher tissue quality in the tumor samples (**Supplementary Table 4**). We note that the total number of transcripts from the MERSCOPE normal TMA run was below what would be typically thought of as a successful run, even when rerunning with the breast panel at 10 µm imaging depth. Such a sample would normally be excluded from analysis, but we continued the data through to illustrate how low transcript capture affects downstream results.

### 10x Xenium shows higher transcript counts per gene without sacrificing specificity

We next sought to directly compare the performance of each iST platform on matched genes. We began with a pseudo-bulk-based approach at the core level since this would not depend on differences in cell segmentation performance (see **Methods**) [21].

First, we examined the run-to-run reproducibility within a single platform for Xenium and MERSCOPE, finding that the total transcript count of all shared genes was highly correlated across data sets acquired with different panels, regardless of the tissue of origin (**Supplementary Fig. 1a**). We also examined the pseudo-bulk gene expression correlation for cores from the same patient in the same dataset and found that correlation was high (Pearson’s r => 0.7) in almost all cases (**Supplementary Fig. 1b-c**), indicating good sample-to-sample reproducibility within a given platform.

To evaluate the relative sensitivity of each platform, we plotted the total transcript counts of every shared gene between all combinations of platform and panel, summed across all matched cores. We found generally linear relationships between all pairs of platforms (**Fig. 2a-c, Supplementary Fig. 2**). Xenium consistently showed higher expression levels on the same genes than CosMx in the tumor TMA, with the Xenium breast having 14.6-fold more counts than the CosMx multi-tissue data sets (**Fig. 2a**). The Xenium multi-tissue panel data showed a slightly smaller difference, with 12.3-fold higher expression on the same genes (**Fig. 2a**), while the lung panel, which was acquired closest in time following slicing, also displayed a median of 14.0-fold higher expression **(Fig. 2a)**. MERSCOPE showed higher expression levels than CosMx when using the breast (10 µm) panel (median of 5.4-fold higher), and comparable expression levels even when using the lung (5 µm) panel (median of 1.1-fold) (**Fig. 2b**). Finally, Xenium showed 2.6-fold higher median expression with the breast panels (10 µm) than MERSCOPE, and 13.6-fold higher median expression with the lung panels (5 µm) **(Fig. 2c)**. In the normal tissue TMA, we found that results were generally consistent, except that the MERSCOPE breast panel showed decreased transcript counts relative to the same panel in the tumor TMA (**Supplementary Fig. 2b-c**), which is consistent with this TMA being unsuccessful for MERSCOPE. Considering the overall higher transcripts per cell across platforms for the tumor TMA **(Supplementary Table 4)**, this suggests that the ability to detect transcripts falls off more strongly with sample quality with MERSCOPE, altering the performance relative to CosMx but not Xenium. Examining the CosMx as compared to Xenium data also revealed an upward curve in the lower expression regime indicative of higher-than-expected calls associated with the low expression regime by CosMx (**Supplementary Fig. 2a**).

**Figure 2:**
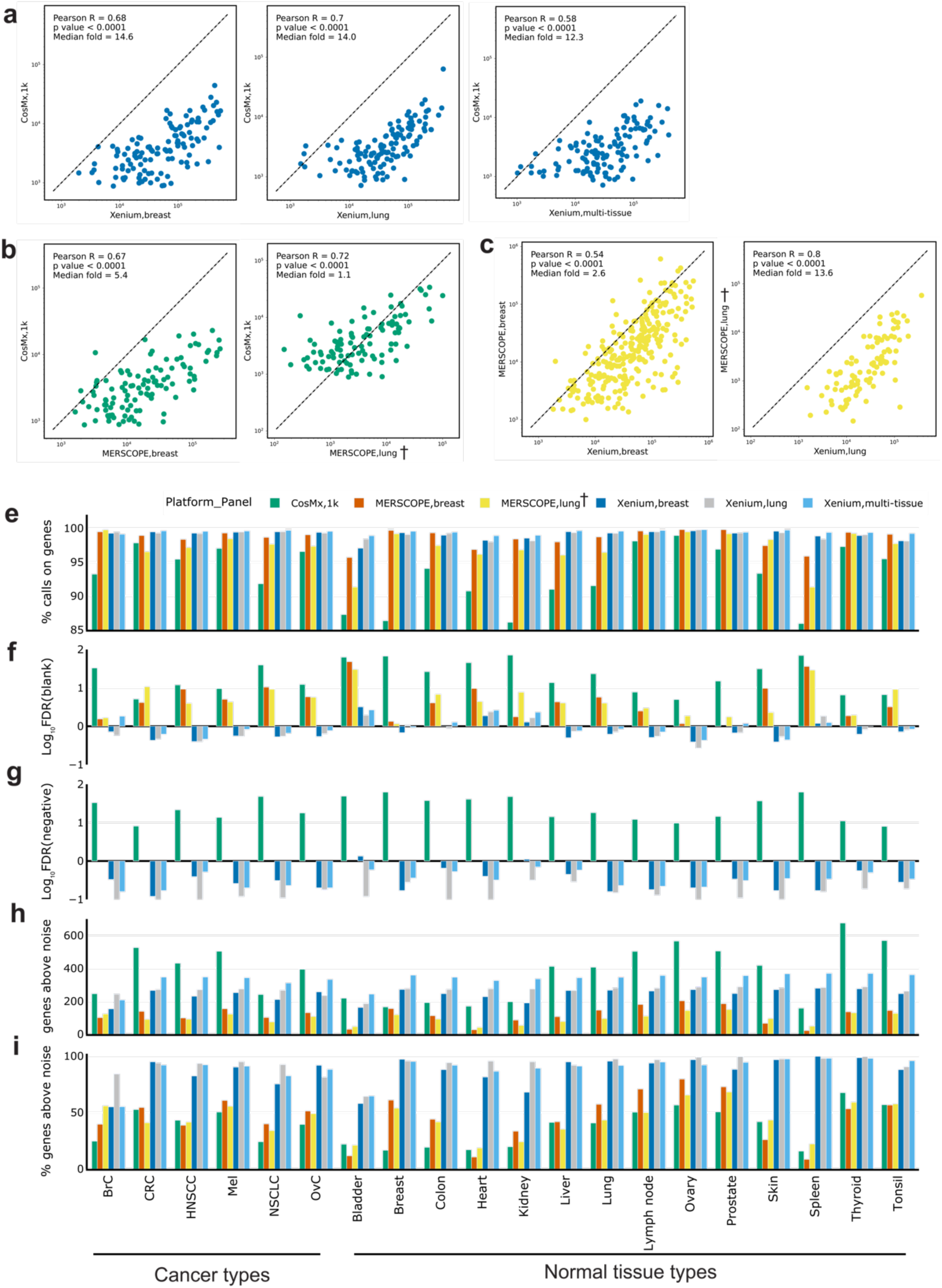
Technical performance comparison of iST platforms grouped by tissue types. **(a)** Scatter plots of summed gene expression levels (on a logarithmic scale) of every shared gene between Xenium (breast/lung) and CosMx (1k) data, captured from matched tumor TMA cores. Each data point corresponds to a gene. **(b)** Same as (a) but between MERSCOPE (breast/lung) and CosMx(1k). **(c)** Same as (a) but between Xenium(breast/lung) and MERSCOPE(breast/lung). **(d)** Same as (a) but between Xenium(multi-tissue) and CosMx(1k). **(e)** Bar plot of percentage of all transcripts corresponding to genes relative to the total number of calls (including negative control probes and unused barcodes) averaged across cores of the same tissue type. Results are presented by panel including breast, lung, and multi-tissue panels from Xenium; breast and lung panels from MERSCOPE; and multi-tissue 1k panel from CosMx. **(f)** Bar plot of false discovery rate (FDR) where FDR(%) = (blank barcode calls / total transcript calls) x (Number of panel genes /Number of blank barcode) x 100. FDR values were log_10_ transformed to better show the differences between panels. **(g)** Same as (f) but using negative control probes to replace blank barcodes. MERSCOPE is missing in this bar plot as it does not have negative control probes by design. **(h)** Bar plot of number of genes detected above noise, estimated as two standard deviations above average of the negative control probes. **(i)** Same as (h) but normalized to the number of genes in a panel. **†** denotes the MERSCOPE lung panel acquired with a 5 µm imaging thickness.

We next wanted to assess the specificity of each platform. Each of the three platforms includes negative controls which are used to evaluate sample quality [22, 23]. Xenium and CosMx include both negative probes (e.g. real probes targeting nucleic acids that are not present in human tissue) and negative barcodes (e.g. algorithmically allowable barcodes that are not associated with any probe in the experimental panel). MERSCOPE includes only negative barcodes by default. To determine specificity, we first calculated the fraction of negative barcodes and probes relative to the number of transcripts for each tissue type **(Fig. 2e).** We found that MERSCOPE and Xenium consistently showed the highest on-target fraction, while CosMx was lower across each tissue type (**Fig. 2e**). However, this measurement is biased because of the relative numbers of controls and target barcodes. We therefore also adopted a false discovery rate (FDR) calculation which normalizes for these differences and is calculated against both the negative probes and negative barcodes (see **Methods, Fig. 2f-g**). We found that Xenium consistently showed the lowest FDR while CosMx showed the highest FDR regardless of whether we standardized to negative control barcodes or probes. This finding is consistent with the upswing in the gene-gene expression plots in **Fig 2a** and **Supplementary Fig. 2a**—as both indicate a higher FDR at the low end of the gene expression range. These results are consistent when visualized across panels **(Fig. 2e-g)**.

Finally, we used the negative control barcodes to evaluate the number of genes reliably detected by each platform in each tissue type. For each core, we calculated the number of genes that were detected two standard deviations above the average expression of the negative control probes. These numbers were then averaged for cores of the same tissue type. Because the CosMx panel was almost three times larger, it yielded a larger absolute number of detected genes in 14 out of 20 tissue types while the Xenium breast panel was higher in the remaining 6 tissue types (**Fig. 2h**, **Supplementary Table 5**). However, Xenium consistently detected the highest fraction of genes in a panel, followed by MERSCOPE and CosMx (**Fig. 2i**).

### iST platforms are all concordant with orthogonal RNA-seq data sets

In the absence of ground truth, it is difficult to evaluate whether a higher number of expressed genes is representative of increased sensitivity to real biology or increased false positive rates. We thus evaluated the correlation of iST data to reference RNA-seq data. We first aggregated pseudo-bulk normal tissue TMA results from all panels of the three platforms and compared them to data from the TCGA program **(see Methods)** [11]. We observed similar correlation coefficients across all gene panels relative to pseudo-bulk RNA-seq expression data (**Fig. 3a-b, Supplementary Table 8**). However, notably, the CosMx data showed a characteristic upswing in the low expression regime, similar to that observed when plotting gene-by-gene expression against MERSCOPE and Xenium (**Fig. 2, Supplementary Fig. 2**).

**Figure 3:**
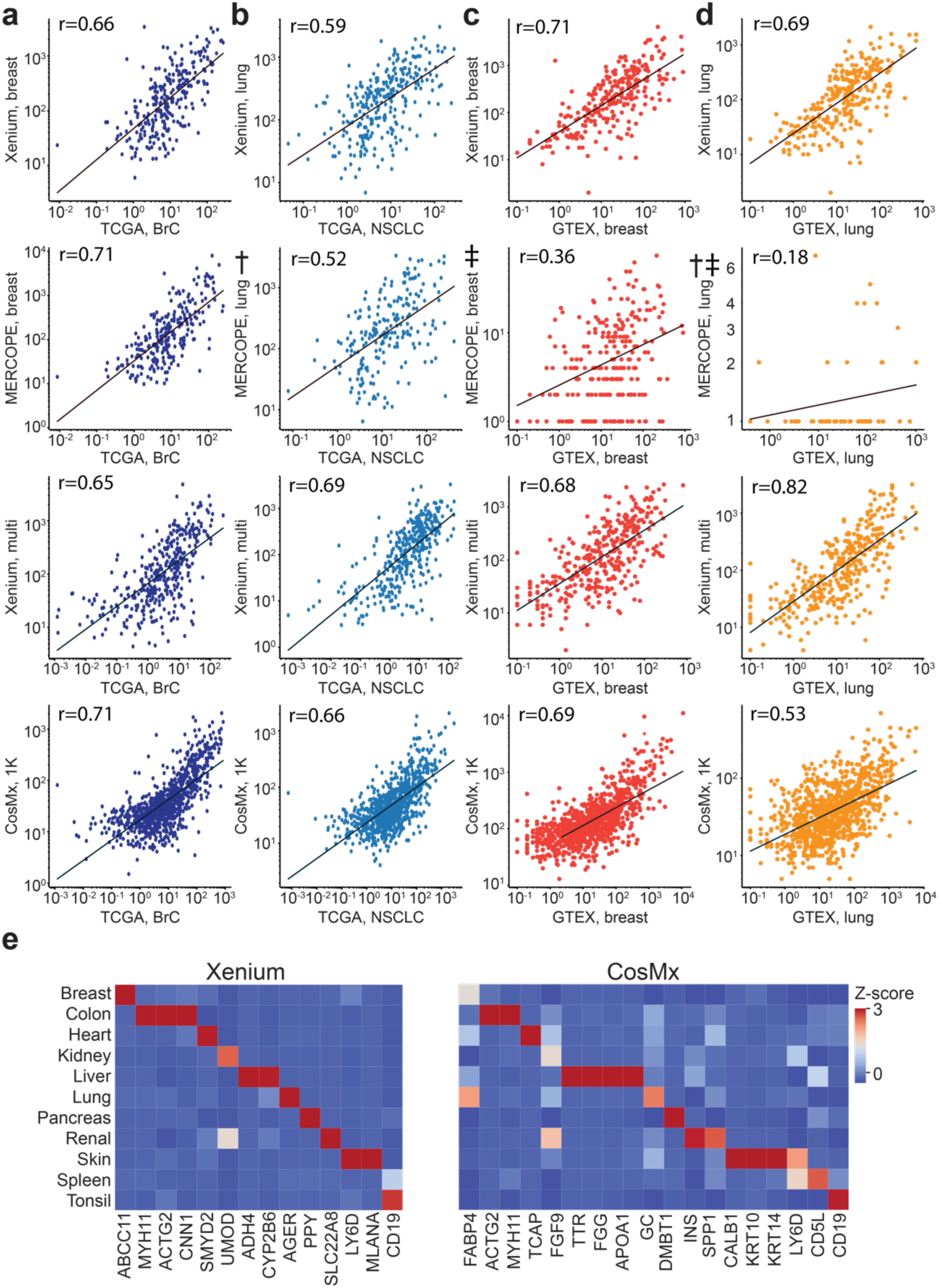
Concordance of iST data with reference RNA-seq datasets. **(a)** Scatter plots of overlapping genes, showing the averaged expression of a gene across breast cancer cores profiled by the indicated panel, normalized to 100,000 vs the average FPKM from TCGA for all samples of a matched tissue type (BRCA). **(b)** Same as (a) but for lung cancer cores plotted vs averaged LUAD and LUSC samples from TCGA. **(c)** Same as (a) but showing breast cores vs averaged nTPM values from GTEx breast samples. **(d)** Same as (a) but for lung cores and samples. † denotes the MERSCOPE lung data acquired with a 5-µm imaging depth on FFPE sample. ‡ denotes the normal tissue TMA data of MERSCOPE which failed initial QC. **(e)** Heatmap of Z-scored average gene expression for several canonical marker genes in the indicated tissue cores for the Xenium multi-tissue panel (left) and CosMx 1K panel (right).

We also compared the pseudo-bulk results from the normal tissue TMA with bulk RNA-seq data obtained from GTEx[12] The Xenium breast, Xenium multi-tissue, and CosMx data sets showed similar correlations to breast data obtained from GTEx, while the MERSCOPE had significantly lower correlation, consistent with a run which doesn’t pass QC (Pearson’s r of 0.36 vs 0.71, 0.68, and 0.69, respectively, **Fig. 3c**). Similar trends were observed in the lung data, with MERSCOPE lung (5 µm) showing the lowest correlation while the other three data sets showed higher correlations to GTEx data (**Fig. 3d**). These relative trends remained true across most normal tissue types, though we found that thyroid, pancreas, and lymph nodes showed the lowest correlations across all panels while prostate, tonsil, and liver showed the highest correlations (**Supplementary Table 9)**. Overall, our comparison to TCGA and GTEx data suggests that while some platforms may be more highly correlated to reference datasets in some cases, all are within a similar correlation regime regardless of tissue type.

We next wanted to determine how the expression of tissue-specific transcript markers varied across each platform. To accomplish this, we curated tissue markers that are unique to each tissue type by selecting genes whose expression in a single tissue exceeds 20 times the sum of other tissues from the GTEx database (see **Methods**). We found tissue-specific expression patterns of several of these markers across all selected panels when visualized across each healthy tissue type (**Fig. 3e**). MERSCOPE showed expression of tissue-specific markers in multiple tissue types, yet some canonical markers were not enriched in certain tissues, potentially caused by the unsuccessful normal TMA data acquisition (Supplementary Fig. 3b). Although CosMx showed satisfying expression patterns for some tissue markers, many canonical markers are not enriched in the expected tissues, possibly due to the high false discovery rate **(Fig. 3f-g)**. Across marker genes, Xenium data had a distinct expression pattern in all tissues, whereas CosMx and MERSCOPE showed a less distinct pattern in many tissue types.

### Out of the box segmentation and filtration can yield cells with comparable numbers of detected transcripts and genes from each platform

Next, we compared the performance of each iST method on a single-cell level. The three platforms generate cell boundaries based on a DAPI image alone (Xenium) or a DAPI image combined with a membrane marker (CosMx and MERSCOPE). When we visually examined the segmentation outputs, Xenium data showed cell boundaries that appeared to include large regions of non-cellular space, in contrast to MERSCOPE and CosMx which tightly followed the visualized cell nucleus (**Fig. 4a**). When transcripts were overlaid with these segmentation boundaries, Xenium cell boundaries fell between regions of transcripts and thus most transcripts were assigned to cells. MERSCOPE and CosMx’s tighter nuclei removed more transcripts, though those that remain appeared more confidently assigned to cells. Overall, when normalized to the imaged tissue area, Xenium and CosMx identified the most putative cells, followed by MERSCOPE (**Fig. 4b, Supplementary Table 6**). In line with the visual inspection, Xenium cells were consistently larger, regardless of data set or panel, followed by CosMx and finally MERSCOPE (**Fig. 4c, Supplementary Table 7**).

**Figure 4:**
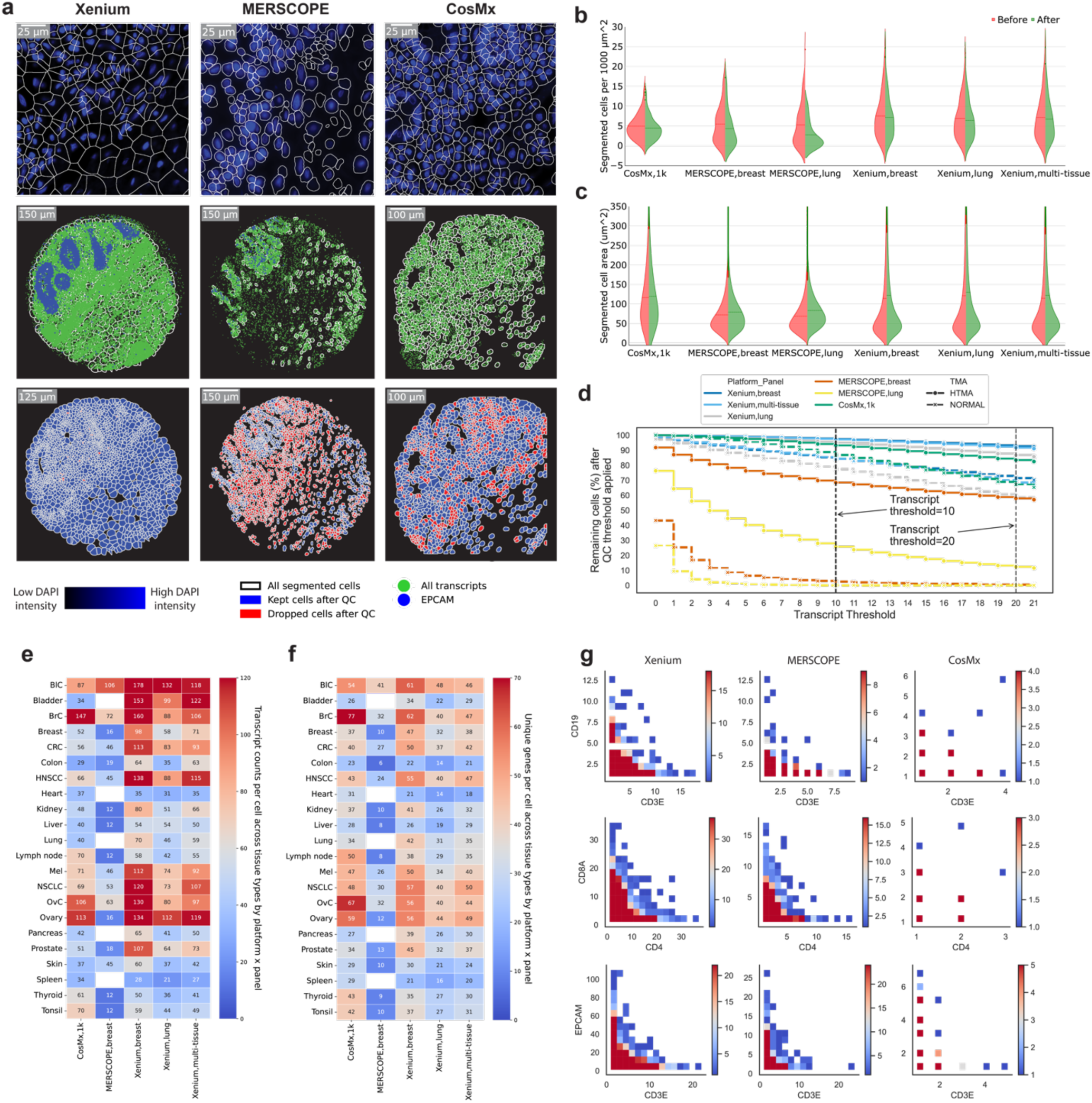
Comparison of cell segmentation results from each iST platform. **(a)** Top row: DAPI image overlaid with cell segmentation boundaries (subset). Middle row: all the transcripts in green dots, white lines for the cell boundaries, and *EPCAM* in blue dots. Bottom row: segmented cell boundaries before and after filtration. **(b)** Violin plot of segmented cells per unit area before (left half) and after filtration (right half) grouped by panel with tumor and normal TMA data combined. **(c)** Same as (b) but showing cell areas before and after filtration. **(d)** Line plot showing remaining cells in percentage after filtering with various thresholds (transcripts per cell). Dotted lines indicate selected thresholds: 10 transcripts or above for Xenium and MERSCOPE and 20 for CosMx. **(e)** Heatmap of transcripts per cell after filtration. All available genes are considered here for each panel. MERSCOPE lung panel (5 µm) was excluded from this heatmap. **(f)** Same as (e) but showing unique genes per cell. **(g)** Co-expression density map for three pairs of disjoint genes (rows) from all three platforms (columns). All cells across all tissues which include at least one detected transcript of either of the indicated genes are plotted together, with color indicating the number of cells at the indicated expression levels of each gene.

We filtered out empty regions of space and cells without any transcripts for downstream examination and quantified the fraction of cells containing differing numbers of transcripts per cell **(Fig. 4d).** We chose a permissive threshold of removing cells with fewer than 10 transcripts for Xenium and MERSCOPE, and 20 transcripts for CosMx from downstream analysis. [11, 24, 25]. The tumor TMA consistently had a greater fraction of cells passing filtration, with Xenium retaining the most cells (97.43% breast, 97.10% multi-tissue, 95.08% lung) followed by CosMx (83.41%) and MERSCOPE (68.46% breast (10 µm), 25.77% lung (5 µm) (**Supplementary Table 3**). The normal tissue TMA had overall lower cell retention performance, but the relative performance of the platforms based on the fractions of cells remained the same. Notably, while CosMx and Xenium still retained > 77% of the cells, MERSCOPE data of the normal tissue TMA had < 3% of cells retained and was thus not used in downstream analysis. Unsurprisingly, filtration decreased the number of retained cells per unit area for all platforms, with the smallest decrease coming for CosMx. The cells retained from CosMx had similar areas, while filtration of the Xenium and MERSCOPE data sets resulted in a higher average cell area (**Fig. 4c**).

After filtration, we compared the number of transcripts and the number of unique genes per retained cell across all tissues and all panels, focusing only on cores that were sampled by all three platforms **(Fig. 4e-f**). Xenium breast panel gave the highest numbers of transcripts per cell in most tissue types, 17 out of 22. The CosMx data showed the highest numbers of transcripts in heart, lymph node, spleen, thyroid, and tonsil; and comparable transcript counts in breast cancer, ovary, and ovarian cancer to the Xenium breast panel. The MERSCOPE data generally had the lowest number of transcripts per cell, though bladder cancer and breast cancer measured with the MERSCOPE breast panels approached the results from Xenium, and the bladder cancer and skin data sets had higher transcripts per cell than CosMx. As expected given its larger panel size, CosMx found many unique genes per cell, showing the largest numbers in 9 tissue types: breast cancer, colon, heart, lymph node, ovary, ovarian cancer, spleen, thyroid, and tonsil; while Xenium breast panel found the most unique genes per cell in 12 tissue types: bladder cancer, bladder, breast, CRC, HNSCC, kidney, lung, melanoma, NSCLC, pancreas, prostate, and skin (**Fig. 4f**). If these analyses were restricted to only the shared genes across all panels, numbers were much lower, but Xenium showed higher expression levels and unique numbers of genes than either CosMx or MERSCOPE (**Supplementary Fig 3c-d**).

We then wanted to determine how effectively different iST platforms’ segmentation algorithms perform. We examined the co-expression of *CD19*, a canonical B-cell marker, and *CD3e*, a canonical T-cell marker across all filtered cells; the co-expression of *CD8* and *CD4*, markers of T-cell subsets; and the co-expression of *CD3e* and *EPCAM*, a marker for epithelial cancer cells [26, 27]. All these marker gene pairs are disjointly expressed, and a well-performing segmentation algorithm should yield few cells expressing both markers. We pooled all the filtered cells from matched cores and all available panels of each platform and plotted the expression of one gene against the other and converted the scatter plot to a heatmap to show cell fractions. We found that Xenium—despite its less visually accurate cell boundaries—and MERSCOPE, showed clear patterns of disjoint expression, separating cells from different lineages, while CosMx showed such a pattern for *EPCAM* vs *CD3e* but not for the other two pairs (**Fig. 4g**). Given the low counts of the immune genes, it was difficult to determine if these were false positive calls or segmentation errors. Nevertheless, since the CosMx panel is much higher plex, and retained similar numbers of transcripts and genes to Xenium, we wondered how these two methods performed in terms of cell type recovery.

### Clustering analyses reveal differences in cell type recovery across platforms

In a typical iST workflow, a key step is reducing the dimensionality of the data by identifying cell types, their unique states, and their expression patterns for further analysis leveraging spatial information[28]. To compare across platforms, we clustered the data from the filtered cells from all the cores for each TMA with a focus on breast tissues. The initial clustering of TMAs from datasets (except MERSCOPE normal tissue) showed expected batch effects caused by patients and tissue types with broadly similar cluster arrangements around morphological tissue features (**Supplementary Fig. 4a-d**). We removed batch effects (**see Methods**) and then performed targeted clustering and cell type annotation for breast samples from the CosMx and Xenium breast datasets; lung samples from the CosMx and Xenium lung datasets; and breast cancer from the CosMx, MERSCOPE breast, and Xenium breast datasets.

In breast samples, we were able to identify nine cell types, including all known major cell types, (adipocytes, alveolar cells, B cells, basal cells, fibroblast cells, hormone-sensing cells, myeloid, T cells, vascular & lymphatic cells) from the Xenium data, using previously established markers (**Fig. 5a, Supplementary Fig. 5**) [29,30,31]. In the CosMx data, we were only able to identify six cell types, including several major cell types, but failed to recognize cell subtypes (B cells, basal cells, fibroblast, hormone-sensing cells, immune cells, and vascular & lymphatic cells) (**Fig. 5a, Supplementary Fig. 5**). A high gene-to-gene correlation was found between all overlapping cell types between Xenium and CosMx (**Fig. 5a**). Similarly, in the lung samples, we were able to identify nine cell types (alveolar epithelial type 1 cell, alveolar epithelial type 2 cell, endothelial capillary cells, endothelial cells, fibroblasts, immune cells, macrophages, mast cells, and stroma cells) in the Xenium lung panel, successfully covering all known major cell types (**Fig. 5b, Supplementary Fig. 5**) [32,33]. Four of the six major clusters were identified and annotated in the lung samples from the CosMx data (endothelial cells, epithelial cells, immune cells, and macrophages), while the other two clusters remained difficult to annotate due to the non-traditional enriched gene markers (**Fig. 5b, Supplementary Fig. 5**). Correlation heatmaps show a strong correlation between the two macrophage clusters identified in Xenium and CosMx (**Fig. 5b**). The epithelial cell cluster from CosMx correlates strongly with alveolar epithelial type 1 cell from Xenium and the endothelial cell cluster from CosMx correlates with endothelial capillary cell cluster from Xenium (**Fig. 5b**).

**Figure 5:**
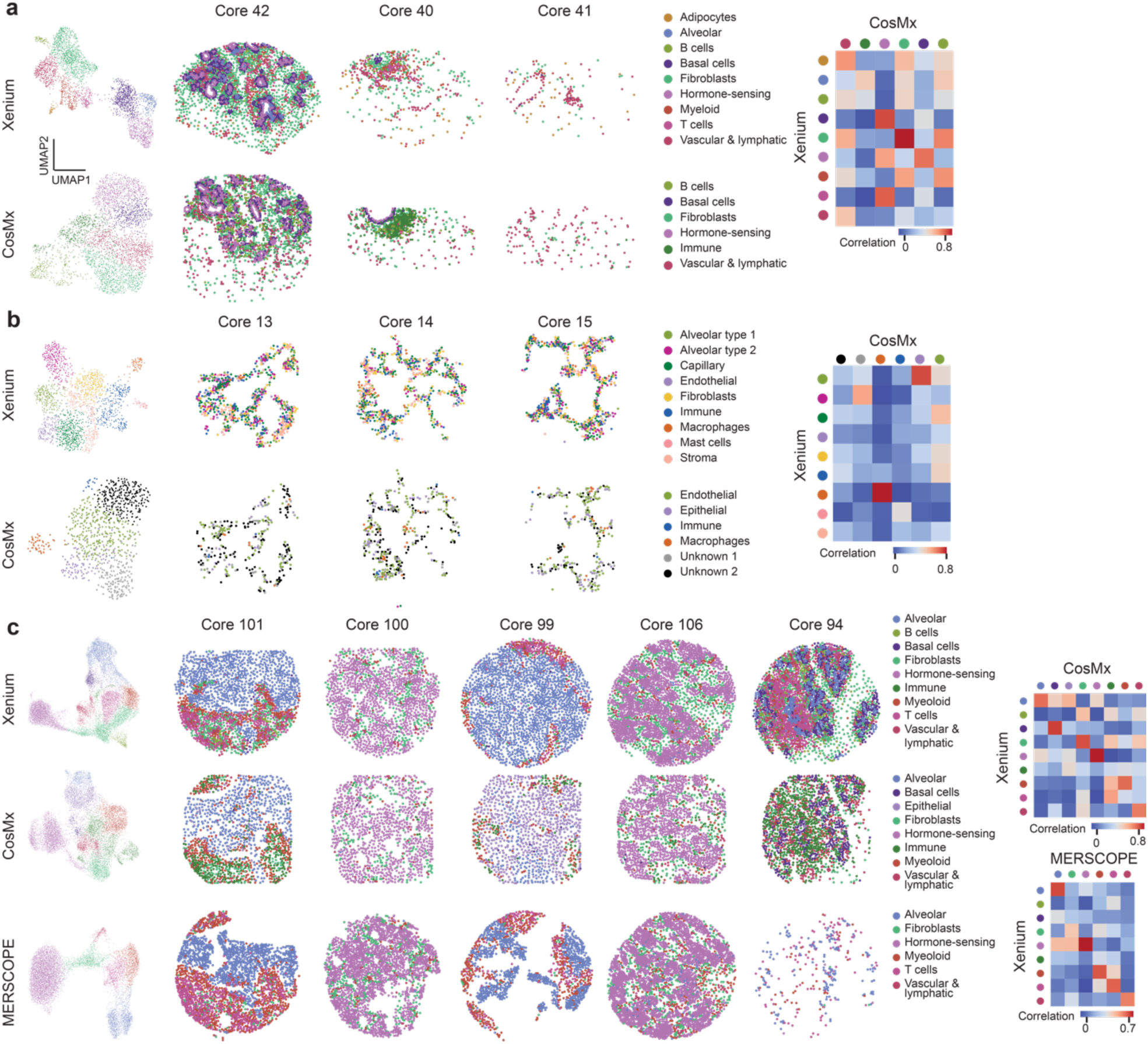
Cell type recovery performance across technology. **(a)** Clustering results of breast samples in normal TMA from Xenium breast panel and CosMx multi-tissue panel. Correlation plot showing the correlation between cell types identified. **(b)** Clustering results of lung samples in normal TMA from Xenium lung panel and CosMx multi-tissue panel. Correlation plot showing the correlation between cell types identified. **(c)** Clustering results of breast cancer samples in tumor TMA from Xenium breast panel, MERFISH breast panel, and CosMx multi-tissue panel. Correlation plot showing the correlation between cell types identified in CosMx and Xenium as well as MERFISH and Xenium.

Finally, in breast cancer, after batch effect removal (**Supplementary Fig. 5d-f**), Xenium resulted in nine cell types (alveolar cells, B cells, basal cells, fibroblast, hormone-sensing cells, immune cells, myeloid, T cells, and vascular & lymphatic cells) (**Fig. 5c, Supplementary Fig. 5**) [34,35,36]. On the other hand, CosMx resulted in eight cell types (alveolar cells, basal cells, epithelial cells, fibroblast cells, hormone-sensing cells, immune cells, myeloid, and vascular & lymphatic cells). MERSCOPE resulted in six cell types, including alveolar cells, fibroblast cells, hormone-sensing cells, myeloid cells, T cells, and vascular & lymphatic cells. The cell type annotation of Xenium and CosMx is comparable in terms of both transcriptomic profile and subtype depth, with CosMx only unable to annotate immune cell subtypes (B cell and T cell). Gene expression of the same cell type from both platforms correlated well (**Fig. 5c, Supplementary 5**). The cell type annotation of CosMx, however, was especially difficult compared to Xenium because of its atypical gene markers shown for each cluster in the heatmaps (**Supplementary Fig. 5**) and low expression of transcripts from canonical markers (**Supplementary Fig. 5g-h**). MERSCOPE, on the other hand, identified most, but not all, the cell types recognized by Xenium and CosMx, including alveolar cells, fibroblast cells, hormone-sensing cells, myeloid, T cells, and vascular & lymphatic cells. MERSCOPE and Xenium showed a high correlation for almost all matching clusters. The correlation map shows a clearer one-to-one mapping between MERSCOPE and Xenium clusters than Xenium and CosMx clusters.

## DISCUSSION

In this study, we compared data obtained with three commercially available iST platforms with archival FFPE tissues to assess overall technical performance and help guide experimental design with human samples that represent an important use case of these platforms. We focused our analyses on technical performance as a function of tissue type, including 7 different tumor types and 16 normal tissue types. Overall, we found that each iST platform presented various tradeoffs in terms of implementation, panel design and panel options, and resulting total transcript quantification and downstream analyses, including cell segmentation, cell quality, and biological interpretation. All these factors must be considered when designing iST experiments.

There are significant workflow differences between the different platforms which factor into the choice of method. Cutting samples onto MERSCOPE coverslip is more difficult than on standard microscope slides. The total hands-on time for running a slide on Xenium is 2-3 days compared to 5-7 days for MERSCOPE and 2 days on CosMx. We found that MERSCOPE and CosMx are well set up for batch processing in the wet lab, either due to built-in pause points or the instrument’s ability to run multiple samples. Xenium is limited for batch processing by a need for a separate thermocycler for each slide pair processed in parallel. After staining, selecting regions of interest (ROIs) presented a surprising challenge for some systems: the Xenium platform could readily image the entire slide as a single ROI which easily covered entire TMAs, but the MERSCOPE ran into a 1cm^2^ imaging area limit which meant cores in the addressable region were left unimaged, while the CosMx workflow required a demanding manual selection of ROIs for each core. These factors are likely to change as each company updates its protocol, but currently, Xenium offers the shortest and least hands-on workflow.

From a technical perspective, we analyzed each resulting dataset with a combination of manufacturer recommended processes for each platform and computational tools that can be implemented by the user downstream. These pipelines each result in count matrices and detected transcripts that can be analyzed using a whole suite of emerging tools. For our purposes, when analyzed at a core level to abrogate the effects of individual cell-segmentation performance, we found that the total number of transcripts varied substantially across iST platform, with Xenium yielding the highest number of transcripts captured followed by CosMx. Indeed, this trend held when normalized for the number of cores imaged and on a per cell basis per area. When this analysis was also restricted to shared genes, we also found that Xenium consistently had higher expression levels across each tissue type, with no clear differences between performance on either tumor or normal tissue.

Using a pseudo-bulk approach, again at the core level, we assessed overall correlation, reproducibility, and sensitivity of each platform. We found high correlation between replicates of the same patient, suggesting that there is high reproducibility across technical replicates on each platform. This is important to consider since cost or input material availability can be prohibitive to implementing experimental designs that leverage technical replicates—though additional tissue may still be valuable for powering cell-cell interaction analysis. We additionally found high correlation on a gene-by-gene basis between MERSCOPE and Xenium platforms. Xenium and MERSCOPE also showed consistently high specificity across tissue types. CosMx displayed a characteristic upward curve when compared to MERSCOPE or Xenium on a gene-by-gene basis, indicating more frequent calls in the lower expression regime. This, coupled with the lower specificity across several tissues for CosMx and the high false discovery rate, suggest that CosMx is prone to errors in calling lowly expressed genes. Finally, Xenium had the highest sensitivity across tissues. CosMx and MERSCOPE both detected fewer transcripts than Xenium. MERSCOPE outperformed CosMx in the higher quality (as judged by relative performance across all platforms) tumor TMA but underperformed it in lower quality normal tissue TMA. In general, our analyses also suggest similar performance within a given platform across a vast array of tissue types assayed here. We note that given the small number of replicates from each tissue, particularly in the normal tissue, we stop short of making blanket statements about relative performance across a particular tissue type. The data suggests, instead, that Xenium and MERSCOPE provide more reliable true-positive signals of lowly expressed genes and that Xenium’s overall performance is less dependent on sample input quality than the other two platforms. MERSCOPE, especially, appears to be particularly sensitive to sample input, highlighting the importance of prescreening RNA integrity according to manufacturer instructions.

When we compared each dataset to existing RNA-seq datasets, we found comparable correlation of pseudo-bulk data to RNA-seq data from GTEx or the TCGA across each panel and platform. However, the presence of a characteristic upswing for CosMx, even when comparing to orthogonal data, further shows that there is a higher false positive rate for lower expression level genes in CosMx data. This upswing could be explained by the absence of probing genes in a particular tissue in a larger panel. However, the Xenium multi tissue panel also includes genes not expressed in breast and lung but does not show a similar upswing. Thus, a more likely interpretation is that the CosMx is prone to a high FDR at the low expression regime. This could also suggest that the CosMx transcript counts and detected gene numbers may be slightly inflated by false discoveries.

From a tissue-specific expression perspective, Xenium showed a distinct expression pattern of key tissue-markers, whereas CosMx and MERSCOPE did not. Additionally, MERSCOPE and CosMx consistently showed expression of known tissue markers in unexpected tissue types. This could be partly explained by an overall low performance on this particular normal tissue TMA for MERSCOPE due to RNA quality. This performance difference could be problematic for studies that are designed to compare tissue-specific factors. For studies whose main biological variables of interest are within the same tissue, factors like sensitivity, specificity, and panel availability may be a more important guide for iST experimental design.

A significant advantage of spatial transcriptomics data is the ability to map expression in single cells. We compared each platform on a cell-level basis by assessing cell identification and cell clustering. Overall, it appears that the out-of-the-box segmentation from Xenium performs poorly in terms of drawing cell boundaries specific to a single cell, while MERSCOPE and CosMx much more closely match cell boundaries. This did not appear to differ on a tissue-by-tissue basis, thus, is likely inherent to the overall approach used by each platform. After applying an expression level filter, Xenium overall retained the highest number of cells across various filtering stringencies. Despite Xenium’s cell boundaries not clearly matching nuclei, both it and MERSCOPE were able to effectively separate cells from different lineage markers, as judged by finding minimal coexpression of disjoint markers, while CosMx showed more double positive cells (out of, it should be noted, fewer cells expressing the target genes overall).

To determine whether clearer identification of lineage markers resulted in improved ability to identify cell types, we performed clustering analyses specifically in the breast tissue and breast cancer samples. We note that we used the full panel, not only the shared genes, when performing these clustering analyses. Xenium allowed for identification of all major cell lineages in the breast when compared to several reference breast atlases. Both the global and tissue clustering results show that CosMx is also able to recognize the major cell types, but cannot identify cell subtypes. Additionally, since the cluster-enriched genes do not correspond to well-known markers, probably due to the low expression caused by low sensitivity and specificity, cell type annotation was particularly difficult. Lastly, despite lower transcript counts and fewer cells, MERFISH still successfully identified cell groups, capturing the patterns seen in other platforms. These differences in cell typing, can also be attributed to the differential performance of cell segmentation pipelines [37,38 39,40]. Since all platforms provide the underlying DAPI stain and morphology images (in the case of CosMx and MERSCOPE) it is likely that segmentation performance could be improved on a sample-by-sample or tissue-type-by tissue-type basis. Future work should seek to assess cell segmentation tools and their performance across data from each platform to help inform the choice of analytical method where needed.

Plex is an important factor in ST experiments which we have not explicitly considered. The kinds of questions that may be answered by a 1,000-plex panel are clearly different than those answered by a 300-plex panel, offering more opportunities to explore intra- and intercellular signaling interactions. Thus, we note that for the right question, the higher false positive rates and lower sensitivities of CosMx relative to Xenium could be tolerated for a broader coverage of the biology. On the other hand, the fully configurable nature of Xenium and MERSCOPE panels could be better suited for branches of biology not well sampled by the 1,000 plex CosMx panel. We recommend subsampling existing atlas data to determine whether the gene set which can be studied will be sufficient to cluster the cell types of interest and identify the necessary biological programs. We note that each of the manufacturers has publicly stated plans to grow their product offerings to increasing panel sizes.

There are several limitations of our study. While we attempted to match time post slicing, the unintentional acquisition of MERSCOPE tissues at thinner thicknesses meant that the rerun MERSCOPE data had a longer time on the slide than other panels. However, the fact that the increase in counts in the rerun matched the increase in imaging volume, and the fact that Xenium runs showed stable expression levels over time suggests that this contribution was minimal. Additionally, our panel design for MERSCOPE required the removal of genes so the panel was compatible with all tissues, lowering the plexity slightly. This could have compromised MERSCOPE’s ability to identify cell types relative to Xenium.

Most importantly, we only attempted to compare the performance of iST platforms under typical use cases for clinical samples obtained from archival biobanks. Our results don’t necessarily extend to non-human samples, frozen samples, and even FFPE samples which have been extensively validated for high RNA integrity. Indeed, there have been reports that MERSCOPE, in previous studies of the mouse brain, shows comparable or even superior results to those reported by 10x Xenium[41]. Given the large change in data quality between the normal and tumor TMA, we cannot exclude the possibility that in the highest quality samples MERSCOPE would provide higher transcript numbers, with the associated downstream benefits relative to Xenium and CosMx. However, the current guidance of DV_200_ > 0.6 restricts studies to the upper regime sample quality and limits archival investigations.

Despite these limitations, our overall interpretation of these results is that amplification of RNA signal is especially important for recovery of transcript counts by iST in low-quality samples where RNA may be highly degraded and fewer landing sites are available for probes. Platforms (such as Xenium) which rely on small numbers of landing sites and are subsequently heavily amplified are robust to RNA degradation and are thus more broadly compatible with a broad range of samples. On the other hand, when sample quality is high (as in some of our tumor samples) the gap between amplified and unamplified platforms’ performance closes and most platforms can yield useful data for subsequent downstream spatial analysis.

## Methods

### Sample choice and TMA construction

Two TMAs were constructed using FFPE clinical discards at Brigham and Women’s Hospital Pathology Core and were acquired with a waiver of consent for non-sequencing based readouts under IRB 2014 P 001721. The samples included:

1. A tumor TMA of 170 cores, 0.6 mm in diameter, including a variety of cancer samples and healthy lymphoid tissue as a positive staining control. The TMA samples were selected from samples previously characterized by ImmunoProfile and were selected to encompass both high and low levels of the biomarkers in the ImmunoProfile panel [*CD8*, *PD-1*, *PD-L1*, *Foxp3*, tumor marker (*Cytokeratin*, *Sox10*, or *PAX8*)]. Annotations were performed by KF and SR based on H&E and immunofluorescence staining. Cores included both tumor and healthy control annotation, though for the purpose of this study, all were combined under their tumor label. Tumors were also chosen to be a mixture of *PD-L1* high and *PD-L1* low, a parameter to be analyzed at a future date. This TMA had previously been studied by both H&E, and several highly multiplexed immunostaining approaches, and was known to be of high morphological integrity.
2. A normal TMA of 45 cores 1.2 mm in diameter representing a broad range of normal tissues. Samples were sourced from the same patient in either duplicate or triplicate.

This TMA was chosen for the breadth of tissue lineages included and the relatively large core size.

All samples were fully de-identified before assembly into TMAs. The breakdown of the number of samples per tissue and the number of cores per tissue is included in **Supplementary Table 1-2**.

### Preparation of sequential sections

Sequential sections were prepared according to manufacturer instructions (“Tissue Preparation Guide Demonstrated Protocol CG000578” for Xenium, “91600112 MERSCOPE User Guide Formalin-Fixed Paraffin-Embedded Tissue Sample Preparation RevB” for Vizgen, and “MAN-10159-01CosMx SMI Manual Slide Preparation Manual” for CosMx) at the Brigham and Women’s Hospital Pathology Core. Prior to collecting samples, ∼50 µm of each TMA were faced off to reach deeper into the sample where RNA integrity was likely higher. 5 µm sequential sections were then collected, floated in a 37°C water bath, and adhered to Xenium slides (10x, PN 1000460), Vizgen FFPE coverslips (Vizgen, PN 10500102), or standard Superfrost+ slides for CosMx (Leica BOND PLUS slides, Leica Biosystems S21.2113.A). TMAs were sliced as close to the center of the active area as possible for each platform. Samples were baked at 42°C for 3 hours for Xenium, 55 °C for 15 minutes for MERSCOPE, and 60°C for 16 hours for CosMx. Sections were stored according to manufacturer instructions prior to processing, with 10x Xenium stored in a desicator at room temperature, Vizgen MERSCOPE coverslips stored at -20°C, and NanoString CosMx slides stored at 4°C. Samples for 10x Xenium and Vizgen MERSCOPE were brought to the Spatial Technology Platform at the Broad Institute for processing, while samples for NanoString CosMx were processed at the Wei lab at Brigham and Women’s Hospital.

### Vizgen MERSCOPE probe selection

Pre-designed probe panels from Vizgen were not available at the time of the experiment. Therefore, we ordered custom gene panels to match the pre-released gene panels from 10x for the human breast and human lung panels. Gene lists were uploaded to the Vizgen panel design portal and were checked against all profiled tissues, removing genes that were overexpressed in any individual tissue based on Vizgen’s design guidelines (FPKM > 900), and ensuring that the total panel FPKM did not exceed the allowed limit in any individual sample type. Panels were manufactured at the 300 gene scale as custom panels BP0892 and BP0893. The final gene lists, for all three iST modalities are available in **Supplementary Table 3.**

### Vizgen MERSCOPE data acquisition

MERSCOPE samples were imaged according to manufacturer protocol “9160001 MERSCOPE Instrument User Guide RevF”. Samples were processed in two batches, the first of four samples, two of each TMA and with each library prepped in parallel; and a follow up sample of each TMA re-run with the breast panel. Samples were first hybridized with anchoring probes overnight before being embedded in a polyacrylamide gel. Samples were incubated for two hours with a digestion solution at 37°C and then overnight at 47°C overnight in a detergent clearing solution and proteinase K to remove native proteins while the anchoring probes kept nucleic acids bound to the gel. After clearing, samples were additionally photobleached using Vizgen’s MERSCOPE Photobleacher for three hours at room temperature in the clearing solution. Samples were hybridized with encoding probes and a cell boundary stain (PN 10400118) and then imaged with imaging kits (PN 10400005). Samples were stored at 37°C in clearing solution after hybridization and before final imaging. After an initial examination of the data, a second batch of both TMAs was run a second time with the human breast panel, increasing the set imaging capture thickness from 5 µm to 10 µm to capture more tissue from cores that had lifted during the gel embedding process. Data was processed on premises through the standard Vizgen workflow to generate cell by gene and transcript by location matrices. We segmented the data with a built-in Cellpose method on the most accurate looking cell boundary stain.

### 10x Xenium data acquisition

10x Xenium samples were processed in three batches according to manufacturer protocols “Probe Hybridization, Ligation & Amplification, User Guide CG0000582” and “Decoding & Imaging, User Guide CG000584”. Samples were stained utilizing 10x’s predesigned Human Breast (10x, PN 1000463), Human Multi-Tissue and Cancer (10x, PN 1000626), and Human Lung panels (10x, PN 1000601), as they became available from the manufacturer. Slides for both TMAs were processed in pairs according to which probe library they were receiving. Slides were stained with a Xenium imaging kit according to manufacturer instructions (10x, PN 1000460). Briefly, padlock probes were incubated overnight before rolling circle amplification and native protein autofluorescence was reduced with a chemical autofluorescence quencher. Slides were processed on a 10 Xenium Analyzer, with ROIs selected to cover the entire TMA region. Data was processed on premises through the standard 10x workflow to generate cell by gene and transcript by location matrices.

### NanoString CosMx

NanoString CosMx samples were prepared with one 1000 plex panel. Samples were hybridized with probes and stained with cell markers. Samples were loaded onto the CosMx SMI at the same time for imaging, during which branched fluorescent probes were hybridized onto the samples to amplify the signal above the background.

NanoString CosMx samples were prepared with Human Universal Cell Characterization 1000 Plex Panel (part number 122000157) according to manufacturer protocol “MAN-10159-01 CosMx SMI Manual Slide Preparation Manual”. Firstly, slides were baked at 60℃ overnight for better tissue adherence. After baking, slides were treated sequentially with deparaffinization, target retrieval (15 min at 100℃), permeabilization (3µg/mL proteinase K, 15 min at 40℃), fiducials application, post-fixation, NHS-acetate application and then hybridized with denatured probes from universal panel and default add-on panel. After *in situ* hybridization (18 hours at 37℃), slides were washed and incubated with DAPI (15 min at RT) and marker stain mix (with *PanCK, CD45, CD68* and cell segmentation marker *CD298/B2M*). Slides were washed and loaded onto the CosMx SMI for UV bleaching, imaging, cycling and scanning. Raw images were decoded by default pipeline on Atomx SIP (cloud-based service). Machine: CosMx_0020. Serial Number: INS2301H0020

### Data preprocessing

After data acquisition, the resulting outputs were uploaded to a Google bucket associated with a terra.bio Workspace for distribution and follow on analysis.

To facilitate standardized data formatting and subsequent analytical processes, we built a data ingestion pipeline with the following objectives: a) to grab cell-level and transcript-level data from diverse platforms and normalize the data structure; b) to tag each cell and transcript with essential metadata including tissue type, tumor status, *PD-L1* status, among others (**Supplementary Fig. 6**); and c) to transform the data into various formats tailored to the requirements of particularized analyses. Specifically, to tag the data, core centers in the TMA were manually identified using DAPI images (Xenium) or cell metadata that contains global coordinates (MERSCOPE and CosMx) using QGIS(version:3.16.10-Hannover). Cells or transcripts within a specified radius were then labeled with core metadata via spatial joining (implemented by GeoPandas, version:0.13.0). In instances where cores are in close proximity or when a uniform radius cannot be applied effectively, we manually generated the core boundary masks.

### Reproducibility

To evaluate panel to panel reproducibility we summed the expression level of shared genes between indicated panels (breast vs. multi-tissue and breast vs. lung panels from Xenium and breast vs. lung panels from MERCOPE) over an individual core and plotted all cores present in each panel, before calculating a Pearson’s correlation. The format of the data used is shown in **Supplementary Table 10.** To evaluate core to core reproducibility, the individual gene counts of core 1 were plotted against those of core 2 and a Pearson’s r correlation was calculated.

### On target rates and false-discovery measurements

To compare across panels and platforms, we subset all datasets to include only cores assayed in all runs. The fraction of on-target barcodes was calculated as a percentage of all transcripts corresponding to genes relative to the total number of calls (including negative control probes and unused barcodes or blank barcodes). These measurements were performed on individual cores and averaged across all cores of the same tissue type.

Because the difference in relative numbers of controls and target barcodes across different platforms, we adopted the false discovery rate (FDR) calculation to evaluate the specificity in a more normalized way (**Fig. 2f-g**). We calculated the FDR of platform *p* panel *m* data in tissue *t* using the following equation and cell level data (**see example in Supplementary Table 11)**:

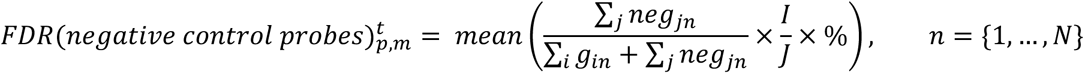

Where *N* is the total number of cores that belong to tissue type *t*, *I* is the total number of unique genes, *J* is the total number of negative control probes, *g_in_* is the gene expression of gene *i* in core *n*, *neg_jn_* is the total calls negative control probe *j* in core.

Since MERSCOPE does not include negative control probes, FDR was recalculated by substituting negative control with blank barcodes (**Fig. 2f**) using the following equation:

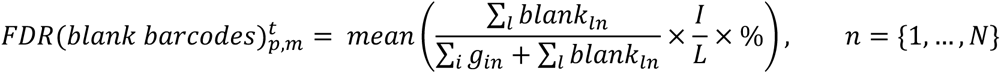

Where *N* is the total number of cores that belong to tissue type *t*, *I* is the total number of unique genes, *L* is the total number of unused barcodes or blank barcodes, *g_in_* is the gene expression of gene *i* in core *n*, *blank_ln_* is the total calls of unused barcode or blank barcode *l* in core *n,* specifically, we used “BLANK” for Xenium, “Blank” for MERSCOPE, and “SystemControl” for CosMx. We only used the data from matched cores, so *N* is same for different platform *p*.

### Sensitivity comparison

Sensitivity was measured by the percentage of the total number of unique genes detected above noise level, where the noise was estimated as two standard deviation above average expression of the negative control probes.

### Orthogonal RNA-Seq concordance analysis

RNA TCGA cancer sample gene data summarizes 7,932 samples from 17 different cancer types, and it provides FPKM for each gene documented. We used all samples which were annotated as BRCA (Breast cancer), BLCA (Bladder cancer), COAD and READ (colorectal cancer), HNSC (head and neck squamous cell carcinoma), LUAD and LUSC (non-small cell lung cancer), SKCM (melanoma), and OV (ovarian cancer). For GTEx, we selected the tissue types matching the annotation in our normal tissue TMA. For each panel, the genes probed by iST were averaged across all patients with the matching tissue label from the RNA-seq database.

To get pseudo-bulked iST values, the expression level of each gene in each core was normalized to the sum of all genes in that core and scaled by 100,000. We then averaged these scaled pseudo-bulk expression values across cores and plotted them against the averaged FPKMs from reference RNA-seq data sets.

### Tissue marker enrichment analysis

To determine the assay’s ability to specifically identify known lineage markers, we focused on the normal tissue TMA profiled with multi-tissue panel of Xenium, breast panel of MERSCOPE, and 1K panel of CosMx. We selected genes with known canonical expression patterns using based on transcriptomics data from GTEx. If a gene had 20-fold higher expression in a specific tissue than every other tissue combined, this gene was considered to be a tissue marker and was used for assessing specificity for each platform. Counts for each gene were normalized to the total counts within the core, and then the Z-score of this gene across tissue types was plotted in a heatmap **Fig. 3e**. We calculated average expression of a gene across cores of the same tissue type and normalized to the total averaged expression of all genes. Z-scores were calculated with the mean and standard deviation across all averaged genes.

### Evaluation of cell segmentation performance

In the absence of ground truth data, we conducted a comparative analysis of cell counts, cell areas, coexpression across various platforms and panels, utilizing the segmentation results supplied by each respective company. To facilitate comparison, cell counts were normalized to a consistent area of 1000 µm². Both cell count and cell area were then delineated at two distinct levels of detail: a consolidated assessment encompassing all tissue types (see **Fig. 4b-c**), as well as a segregated evaluation by individual tissue types (refer to **Supplementary Table 6-7**). To evaluate the biological performance of the segmentation, we plotted coexpression plots of gene pairs that are mutually exclusive including *CD3e* vs. *CD19*, *CD4* vs. *CD8*, and *CD3e* vs. *EPCAM*. We pooled all the filtered cells from matched cores and all available panels of each platform, dropped cells which do not express either gene, plotted the expression of one gene against the other, and converted the scatter plot to a 2D histogram showing cell numbers in each co-expression bin (**Fig. 4g**).

### Cells per area quantification

Segmented cells were aggregated by TMA cores. For Xenium and MERSCOPE data, the estimation of tissue area was performed by calculating the area of a discernible circle, utilizing respective radius of 0.3 µm and 0.6 µm for tumor and normal TMA samples. Conversely, for the CosMx dataset, the tissue area estimation was approached differently due to its square-like data presentation, a result of the FOV selection process. Here, the tissue area was deduced by multiplying the number of FOVs covered by each core with the area of a single FOV.

### Clustering

For cell filtering, cells with less than 10 transcript counts in MERFISH and Xenium datasets were removed, and cells with less than 20 transcript counts in CosMx datasets were removed. We followed standard processes to then cluster and annotate cell types across each dataset using Scanpy[42]. Briefly, data was normalized and scaled, dimensionality reduction was performed and cell clusters were identified[43, 44]. To identify the cell type for each cluster, we used a t-test to find the markers for each Leiden cluster and annotated them according to previous literature[29–36]. These are some of the example markers used for cell type annotation: in breast samples, PIGR and KIT for alveolar cells, for B cells, KRT5, DST, and MYLK for basal cells, LUM, MMP2 and CXCL12 for fibroblast, etc. Heatmaps of the top 3 markers for each cluster are drawn for each dataset from all three panels (refer to **Supplementary Figure 5a-c**). For datasets that showed batch effect with patients, Harmony was used to remove this variance[45]. Correlation heatmaps were generated over overlapping genes that exist in both datasets, and the Pearson correlation coefficient was calculated.

## Acknowledgment

We thank Nir Hacohen, Ilya Korsunsky, Roopa Madhu, and Kseniia Anufrieva for helpful discussions, as well as Patricia Rogers and Natan Pierete for assistance with the 10x Xenium data acquisition. We appreciate 10x Genomics, Vizgen, and NanoString Technologies for reviewing data and analyses for quality. This work is supported by a Broad Institute SPARC grant and by a Brigham and Women’s Hospital Department of Medicine - Broad Institution collaborative research Award. K.W. is supported by a NIH-NIAMS K08AR077037, a Burroughs Wellcome Fund Career Awards for Medical Scientists, and a Doris Duke Charitable Foundation Clinical Scientist Development Award. B.A.G is supported in part through the Geisel School of Medicine at Dartmouth’s Center for Quantitative Biology through a grant from the National Institute of General Medical Sciences (NIGMS, P20GM130454) of the NIH.

## Author contributions

Conceptualization: A.Y., S.F., tissue-microarray construction: K.F., K.P., T.B., S.R., pathological annotation: K.P., S.R., gene selection: A.Y., S.F., Xenium and MERSCOPE data acquisition: J.N., CosMx data acquisition: C.G., M.T.,K.W., analysis: H.W., R.H., B.G.,S.F., figure generation: H.W., R.H., B.G.,S.F., writing original draft: H.W., R.H., J.N., B.G., S.F., draft reviewing and editing: H.W., R.H., B.G., S.F., supervision: B.G., S.F., funding acquisition: S.F. K.W.

## Data and Code Availability

Cell-level data and gene-level data will be made available at the time of final publication. All code used in this manuscript for data processing and analysis will be made available on GitHub prior to final publication.

## Declaration of interests

All authors declare that they have no conflicts of interest.

## Supplementary Figures

**Supplementary Figure 1:**
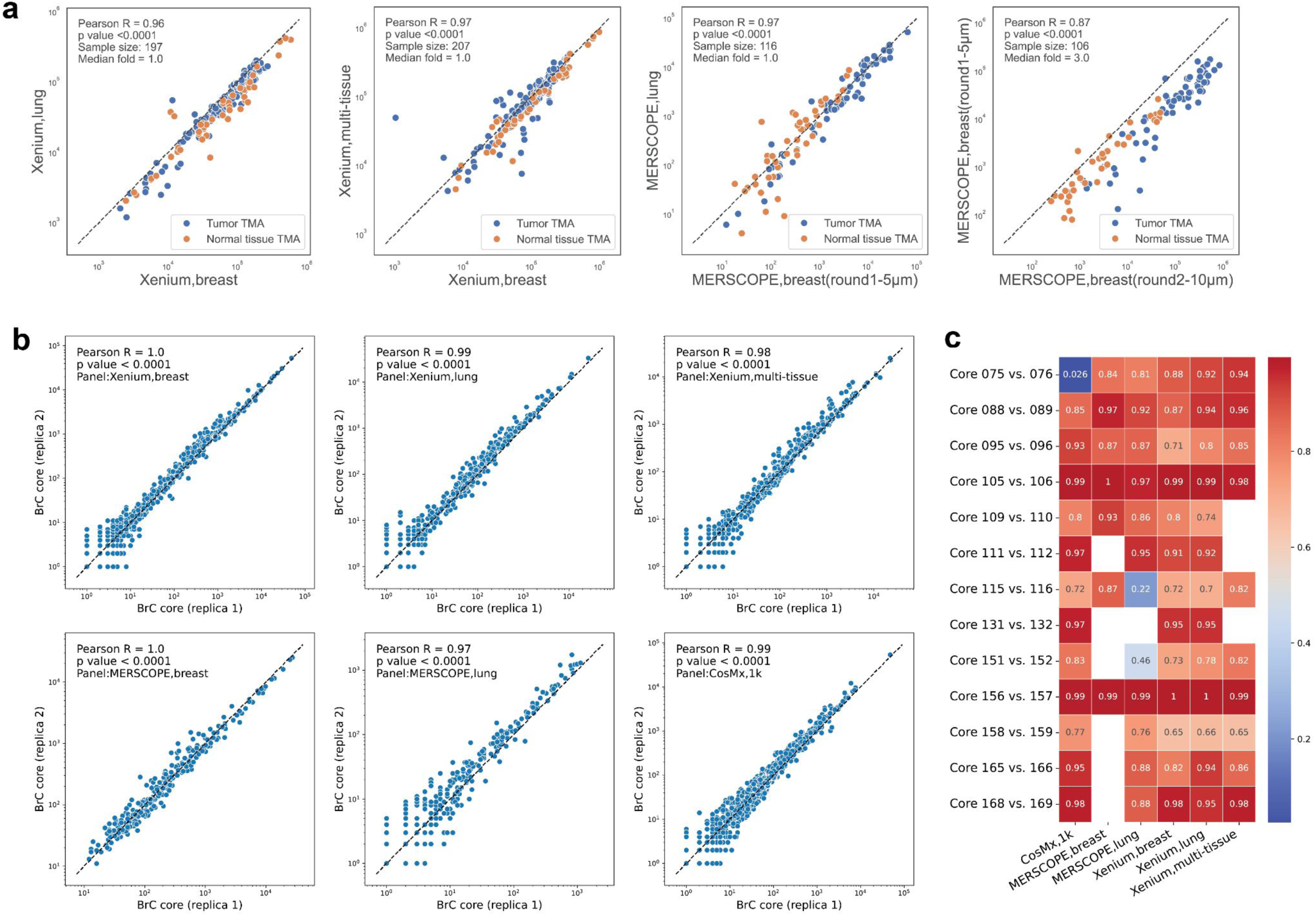
Reproducibility across different panels and cores from same patient. **(a)** Scatter plots of cumulative gene expression levels (on a logarithmic scale) of shared genes between two panels within each platforms, captured from matched tissue cores. Column 1: Xenium breast vs. Xenium lung; Column 2: Xenium breast vs. Xenium multi-tissue; Column 3: MERSCOPE breast round 1(5 µm) vs. MERSCOPE breast round 2(10 µm). Each data point corresponds to a TMA core. **(b)** Scatter plots of gene expression levels (on a logarithmic scale) of every shared gene between two cores of the same tissue type from the same patient. In this example, cores are from breast cancer tissue. Each data point corresponds to a gene. **(c)** Heatmap of correlation coefficient expressed as Pearson’s r values, indicating good core-to-core or sample-to-sample reproducibility. Core pairs are selected from same tissue/tumor type from the same patients.

**Supplementary Figure 2:**
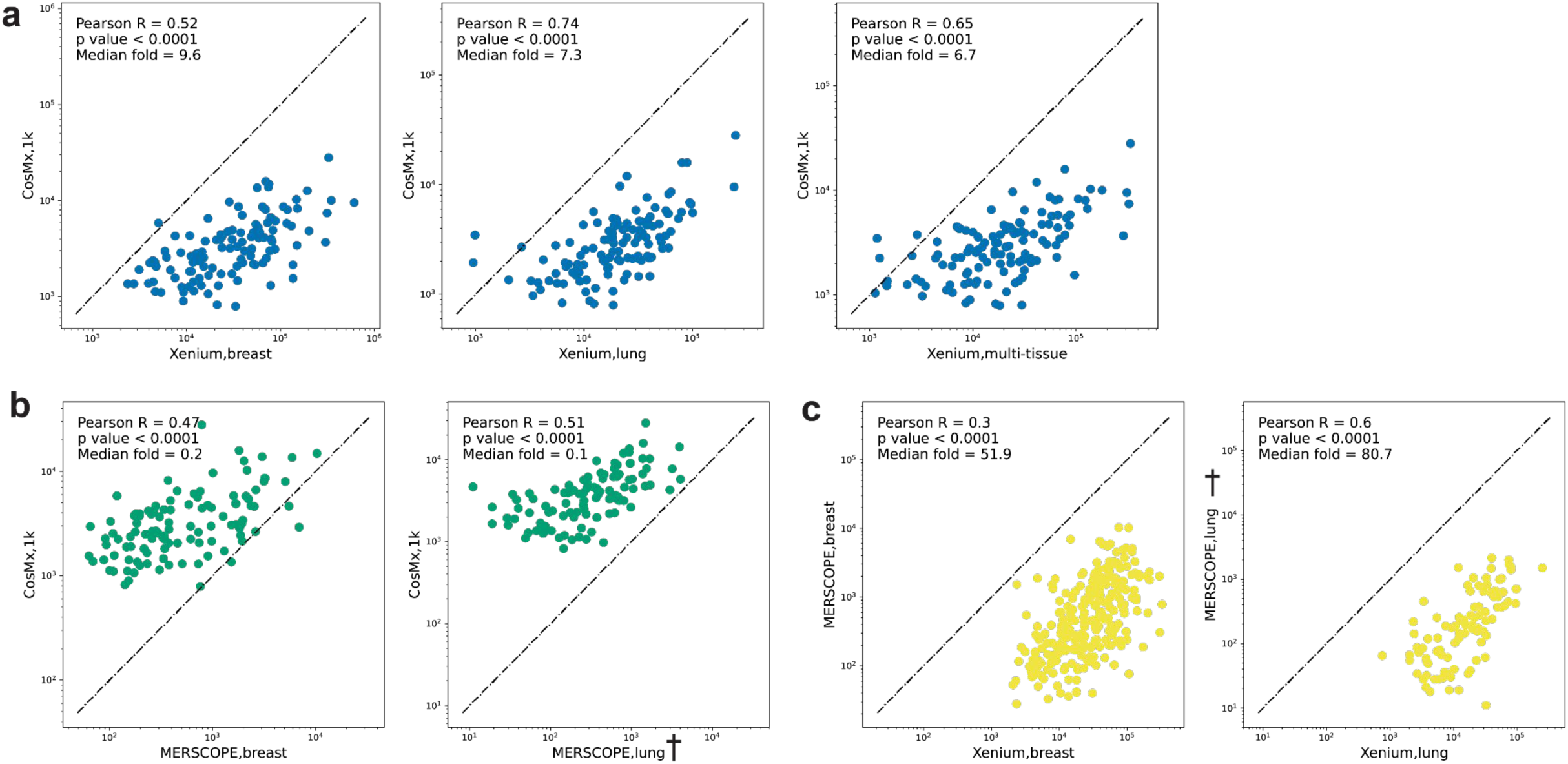
Gene by gene plots of iST results by panel and by tissue microarray. **(a)** Scatter plots of summed gene expression levels (on a logarithmic scale) of every shared gene between Xenium (breast/lung) and CosMx (1k) data, captured from matched normal tissue TMA cores. Each data point corresponds to a gene. **(b)** Same as (a) but between MERSCOPE (breast/lung) and CosMx(1k). **(c)** Same as (a) but between Xenium(breast/lung) and MERSCOPE(breast/lung). **(d)** Same as (a) but between Xenium(multi-tissue) and CosMx(1k). **†** denotes the MERSCOPE lung panel acquired with a 5 µm imaging thickness.

**Supplementary Figure 3:**
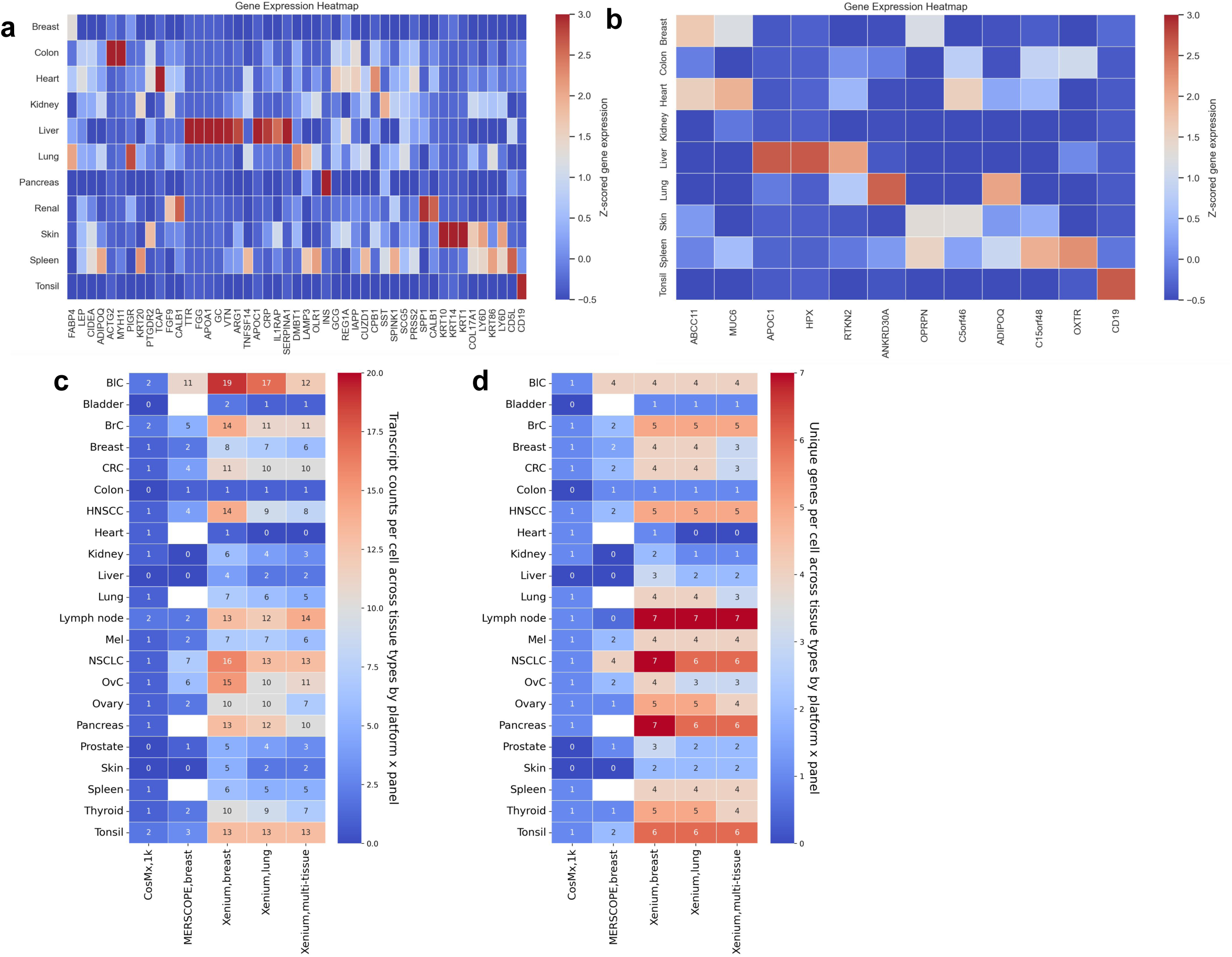
Tissue marker analyses and cell level measurements. **(a)** Heatmap of Z-scored gene expression showing CosMx’s ability to specifically identify known lineage markers. We focused on the normal tissue TMA profiled with multi-tissue panel and selected genes with canonical expression patterns for this analysis. **(b)** Same as (a) but for MERSCOPE (breast panel). **(c)** Heatmap of transcripts per cell after filtration. Only shared genes (40) are considered here for each panel. **(d)** Same as (c) but showing unique transcripts from the same gene set.

**Supplementary Fig: 4.**
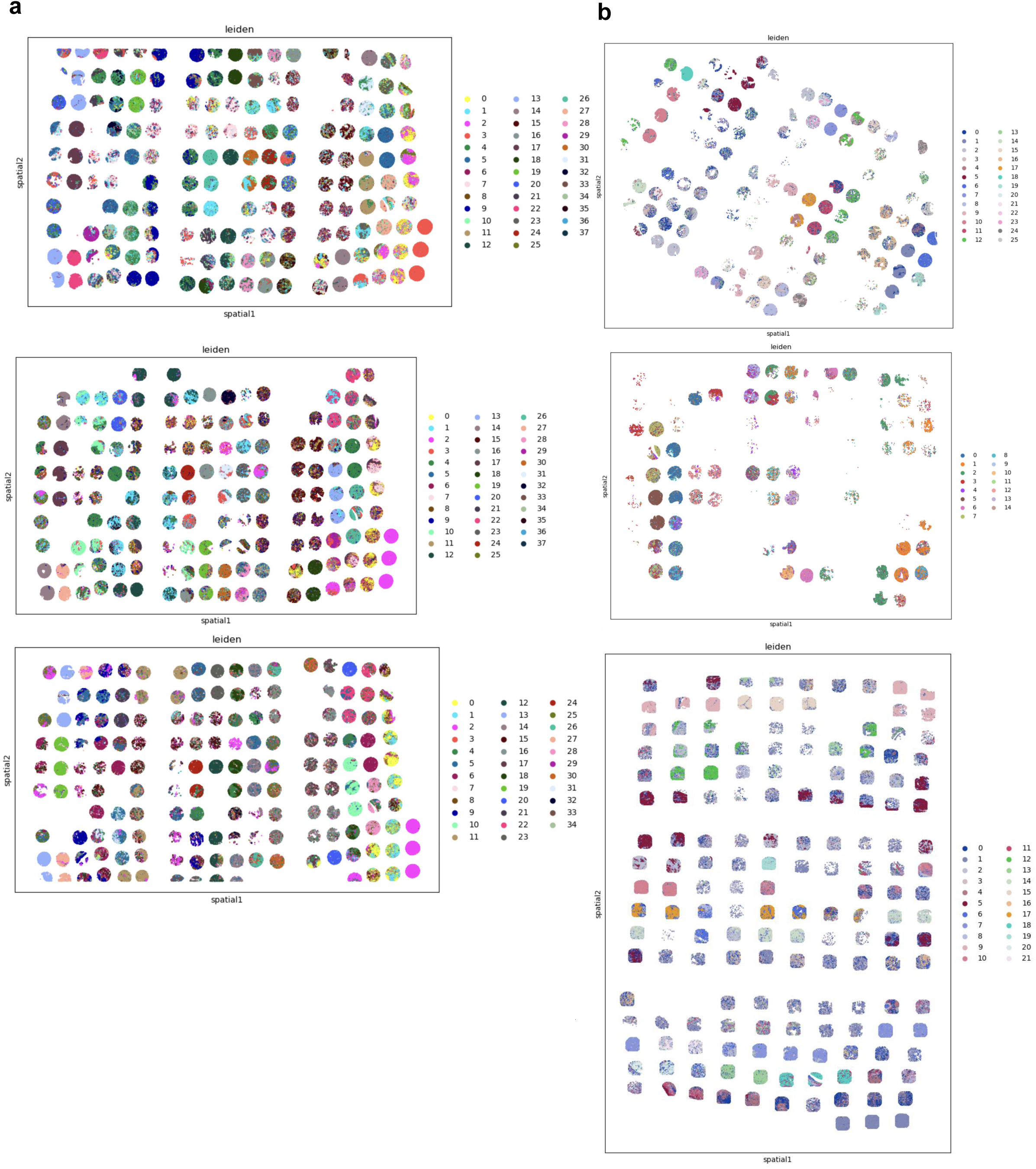
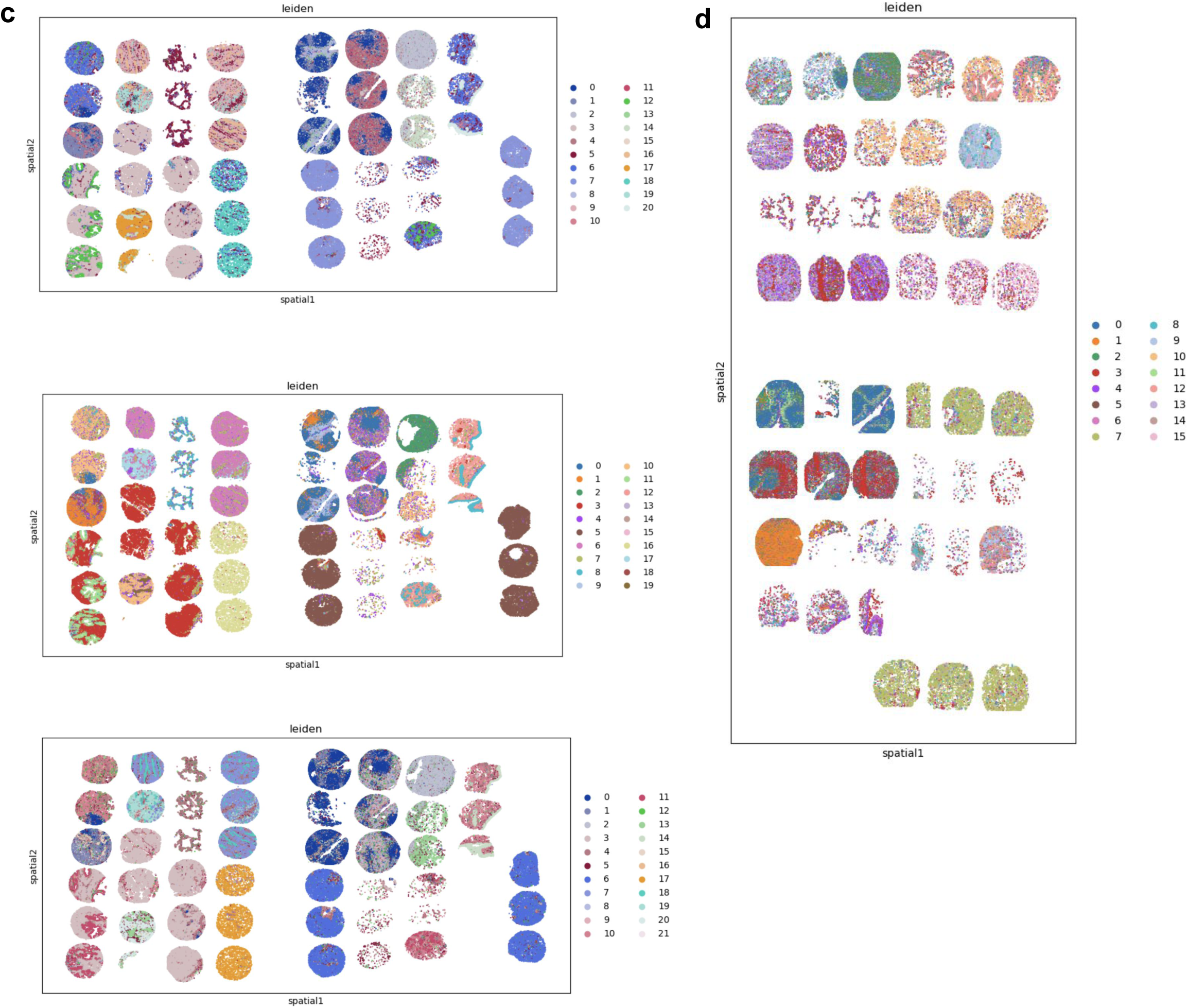
Global clustering analyses. **(a)** Global Clustering results of tumor TMA from Xenium breast panel (top), Xenium lung panel (middle), and Xenium panhuman panel (bottom). **(b)** Global Clustering results of tumor TMA from MERFISH breast panel (top), MERSCOPE lung panel (middle), and CosMx multitissue panel (bottom). **(c)** Global Clustering results of normal TMA from Xenium breast panel (top), Xenium lung panel (middle), and Xenium panhuman panel (bottom). **(d)** Global Clustering results of normal TMA from CosMx multitissue panel (bottom).

**Supplementary Fig: 5.**
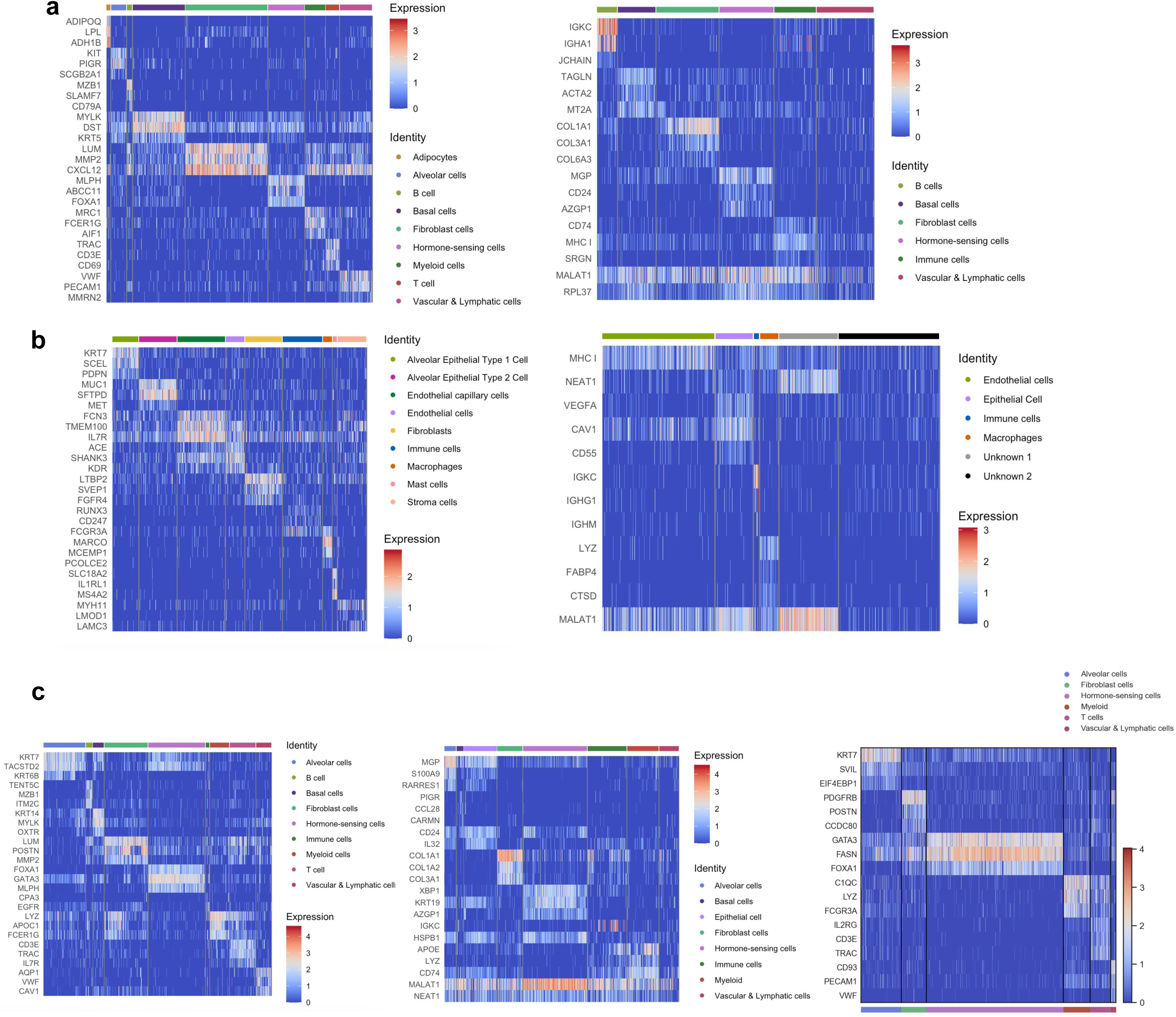
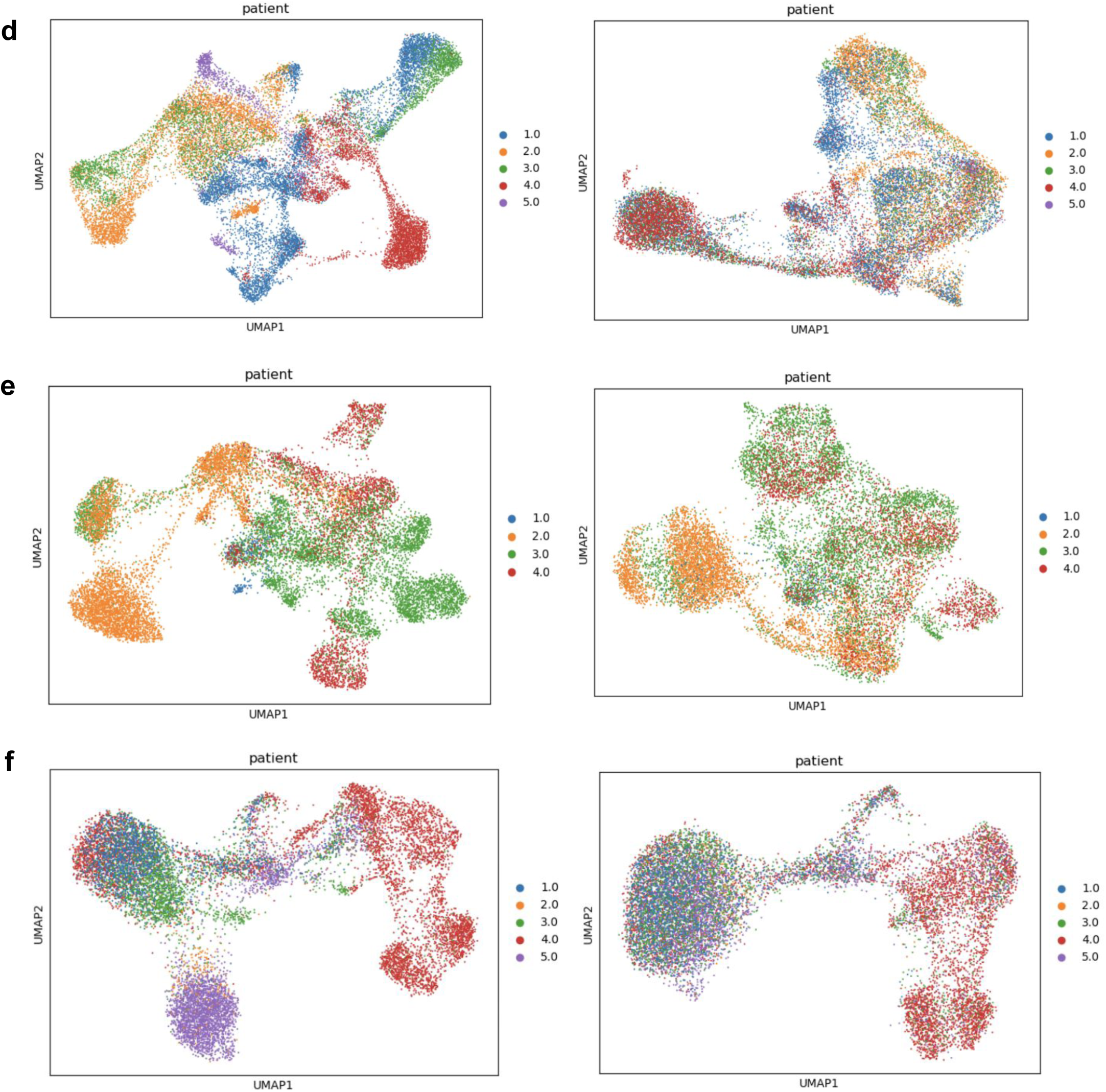
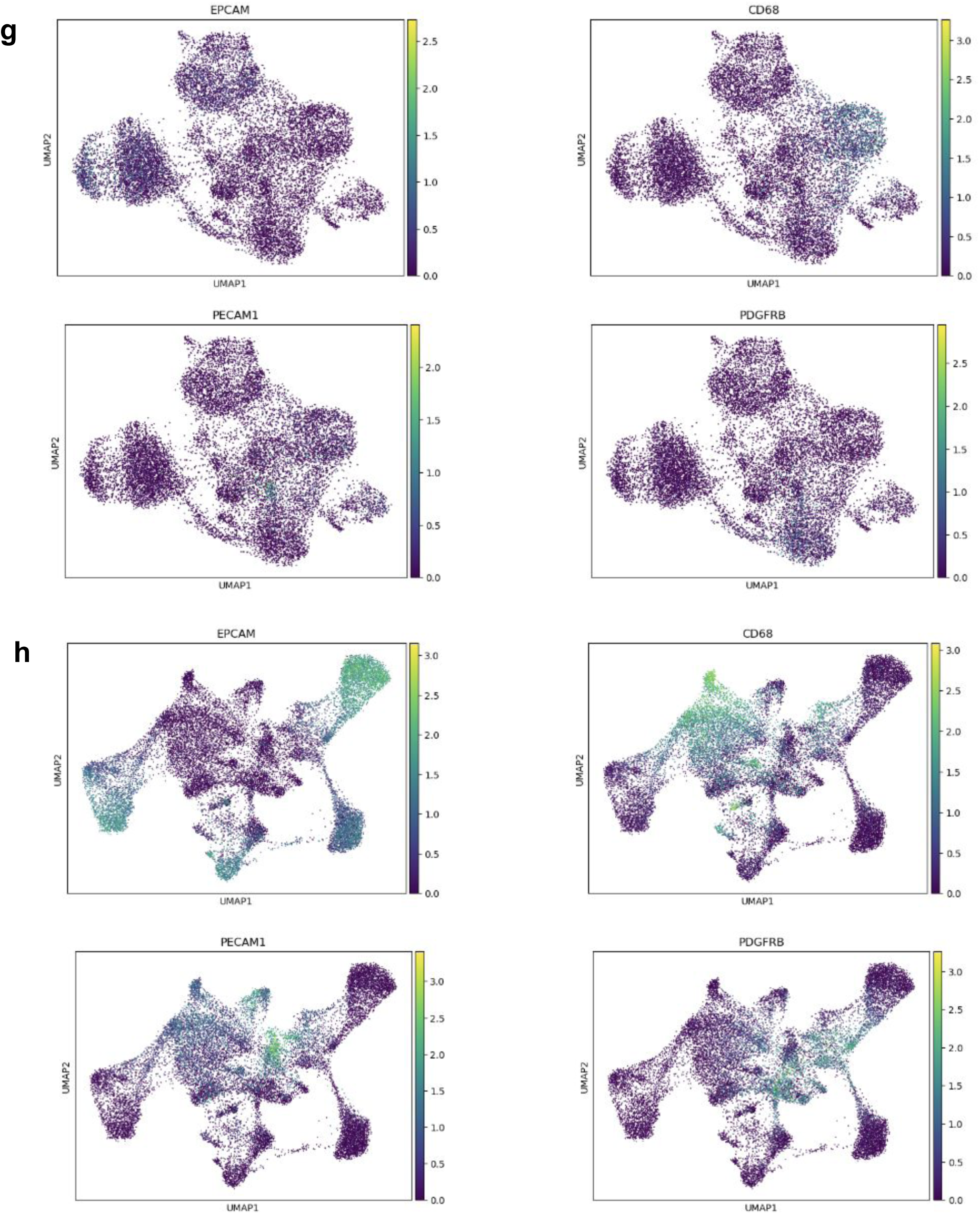
Cell type recovery and UMAPs. **(a)** Heatmap showing the top gene markers for cell types annotated in breast samples of normal TMA from Xenium breast (left) and Cosmx multitissue (right). **(b)** Heatmap showing the top gene markers for cell types annotated in lung samples of normal TMA from Xenium lung (left) and Cosmx multitissue (right). **(c)** Heatmap showing the top gene markers for cell types annotated in breast cancer samples of tumor TMA from Xenium breast (left), Cosmx multitissue (middle), NERSCOPE breast (right). **(d)** UMAP of breast cancer samples of tumor TMA from Xenium breast panel pre (left) and post (right) batch effect removal. **(e)** UMAP of breast cancer samples of tumor TMA from Cosmx multitissue panel pre (left) and post (right) batch effect removal. **(f)** UMAP of breast cancer samples of tumor TMA from MERFISH breast panel pre (left) and post (right) batch effect removal **(g)** UMAP plot of well-known gene markers for BrC, in breast cancer samples of tumor TMA from CosMx multitissue panel. **(h)** UMAP plot of well-known gene markers for BrC, in breast cancer samples of tumor TMA from Xenium multitissue panel.

**Supplementary Figure 6:**
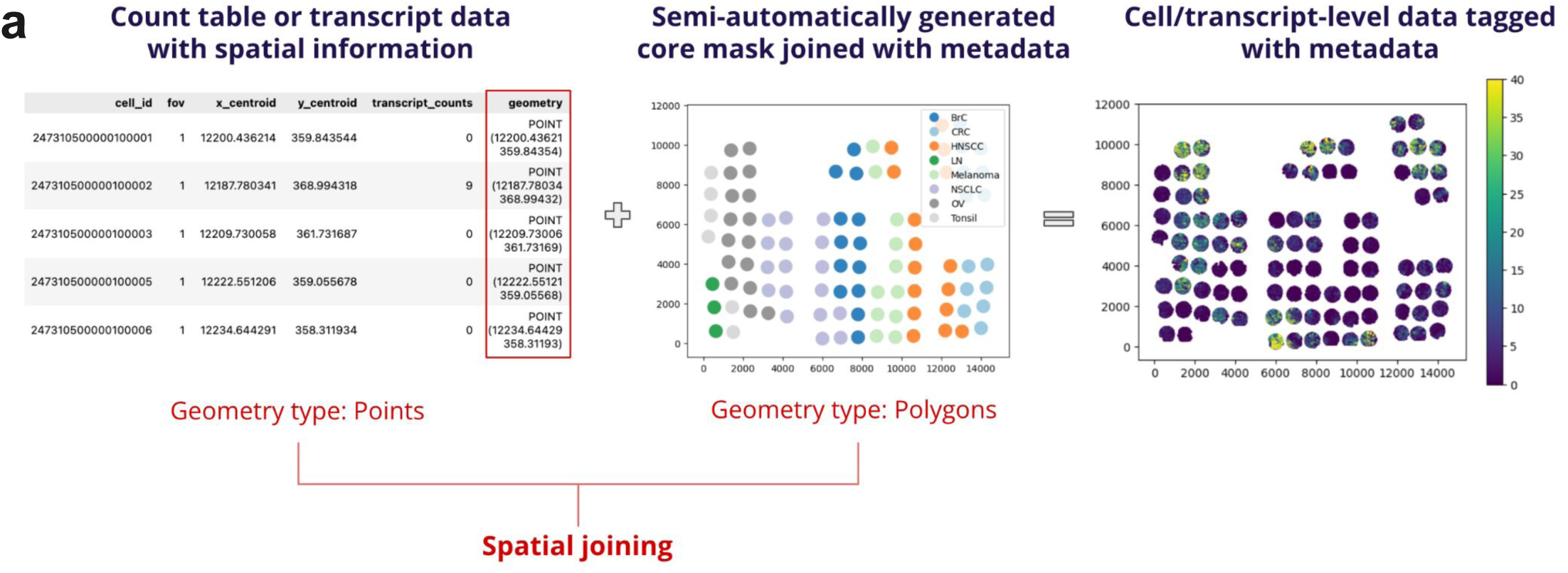
Workflow for tagging imaging spatial transcriptomics data. **(a)** To facilitate standardized data formatting and subsequent analytical processes, we built this data ingestion pipeline with the following objectives: 1) to grab cell-level and transcript-level data from diverse platforms and normalize the data structure; 2) to tag each cell and transcript with essential metadata including tissue type, tumor status, PD-L1 status, among others; and 3) to transform the data into various formats tailored to the requirements of particularized analyses. Specifically, to tag the data, core centers in the TMA were pinpointed using DAPI images (Xenium) or cell metadata that contains global coordinates (MERSCOPE and CosMx) using QGIS(version:3.16.10-Hannover). Cells or transcripts within a specified radius were then labeled with core metadata via spatial joining (implemented by GeoPandas, version:0.13.0). In instances where the cores are in close proximity or when a uniform radius cannot be applied effectively, we manually generated the core boundary masks.

## Notes

https://docs.google.com/spreadsheets/d/14ZChOSscpm24c-BC25M1gu6VR4AEmkAzxsdwFbHYuuo/edit#gid=87987271

### Competing Interest Statement

The authors have declared no competing interest.

### Summary of Updates

We fixed some typos and improved the readability.

