## Supplementary Tables for "Systematic benchmarking of imaging spatial transcriptomics platforms in FFPE tissues"

**Supplementary Table 1:** Tumor tissue-microarray metadata. Abbreviations: bladder cancer (BlC), colorectal cancer (CRC), head and neck squamous cell carcinomas (HNSCC), melanoma (Mel), breast cancer (BrC), non-small cell lung cancer (NSCLC), and ovarian cancer (OvC).

| Core index | Tissue type | Tumor | Patient number | PD-L1 status |
|---|---|---|---|---|
| 1 | BlC | 1 | 1 | high |
| 2 | BlC | 1 | 1 | high |
| 3 | BlC | 1 | 1 | high |
| 4 | BlC | 0 | 1 | low |
| 5 | BlC | 0 | 1 | NaN |
| 6 | BlC | 1 | 2 | high |
| 7 | BlC | 1 | 2 | high |
| 8 | BlC | 1 | 2 | high |
| 9 | BlC | 1 | 3 | NaN |
| 10 | BlC | 1 | 3 | low |
| 11 | BlC | 1 | 5 | low |
| 12 | BlC | 1 | 5 | low |
| 13 | BlC | 0 | 4 | low |
| 14 | BlC | 0 | 4 | low |
| 15 | BlC | 1 | 4 | low |
| 16 | BlC | 1 | 4 | low |
| 17 | BlC | 1 | 4 | low |
| 18 | BlC | 0 | 3 | low |
| 19 | BlC | 0 | 3 | low |
| 20 | BlC | 1 | 3 | low |
| 21 | BlC | 1 | 5 | low |
| 22 | CRC | 1 | 1 | low |
| 23 | CRC | 1 | 1 | low |
| 24 | CRC | 1 | 1 | low |
| 25 | CRC | 0 | 1 | low |
| 26 | CRC | 0 | 1 | low |
| 27 | CRC | 1 | 2 | low |
| 28 | CRC | 1 | 2 | low |

| | | | | |
|---|---|---|---|---|
| 29 | CRC | 1 | 2 | low |
| 30 | CRC | 0 | 2 | low |
| 31 | CRC | 0 | 4 | low |
| 32 | CRC | 1 | 4 | low |
| 33 | CRC | 1 | 4 | low |
| 34 | CRC | 1 | 4 | low |
| 35 | CRC | 0 | 3 | NaN |
| 36 | CRC | 0 | 3 | low |
| 37 | CRC | 1 | 3 | high |
| 38 | CRC | 1 | 3 | low |
| 39 | CRC | 1 | 3 | high |
| 40 | CRC | 0 | 2 | low |
| 41 | CRC | 0 | 4 | high |
| 42 | CRC | 1 | 5 | high |
| 43 | CRC | 1 | 5 | high |
| 44 | CRC | 1 | 5 | high |
| 45 | CRC | 0 | 5 | high |
| 46 | CRC | 0 | 5 | high |
| 47 | CRC | 1 | 6 | high |
| 48 | CRC | 1 | 6 | high |
| 49 | CRC | 1 | 6 | high |
| 50 | HNSCC | 1 | 1 | high |
| 51 | HNSCC | 1 | 3 | high |
| 52 | HNSCC | 0 | 2 | high |
| 53 | HNSCC | 0 | 2 | low |
| 54 | HNSCC | 1 | 2 | low |
| 55 | HNSCC | 1 | 2 | high |
| 56 | HNSCC | 1 | 2 | high |
| 57 | HNSCC | 0 | 1 | high |
| 58 | HNSCC | 0 | 1 | high |
| 59 | HNSCC | 1 | 1 | low |
| 60 | HNSCC | 1 | 1 | high |
| 61 | HNSCC | 1 | 3 | low |
| 62 | HNSCC | 1 | 3 | high |

| | | | | |
|---|---|---|---|---|
| 63 | HNSCC | 0 | 3 | low |
| 64 | HNSCC | 0 | 3 | high |
| 65 | HNSCC | 1 | 4 | low |
| 66 | HNSCC | 1 | 4 | high |
| 67 | HNSCC | 1 | 4 | low |
| 68 | HNSCC | 0 | 4 | low |
| 69 | HNSCC | 0 | 4 | high |
| 70 | HNSCC | 1 | 5 | high |
| 71 | Mel | 1 | 2 | high |
| 72 | Mel | 0 | 1 | low |
| 73 | Mel | 0 | 1 | low |
| 74 | Mel | 1 | 1 | low |
| 75 | Mel | 1 | 1 | low |
| 76 | Mel | 1 | 1 | low |
| 77 | HNSCC | 0 | 5 | NaN |
| 78 | HNSCC | 0 | 5 | low |
| 79 | HNSCC | 1 | 5 | high |
| 80 | HNSCC | 1 | 5 | low |
| 81 | Mel | 1 | 2 | low |
| 82 | Mel | 1 | 2 | high |
| 83 | Mel | 0 | 2 | high |
| 84 | Mel | 0 | 2 | high |
| 85 | Mel | 1 | 3 | high |
| 86 | Mel | 1 | 3 | low |
| 87 | Mel | 1 | 3 | NaN |
| 88 | Mel | 1 | 4 | low |
| 89 | Mel | 1 | 4 | low |
| 90 | Mel | 1 | 4 | low |
| 91 | BrC | 1 | 3 | high |
| 92 | BrC | 1 | 3 | high |
| 93 | BrC | 0 | 2 | high |
| 94 | BrC | 0 | 2 | high |
| 95 | BrC | 1 | 2 | high |
| 96 | BrC | 1 | 2 | high |

| | | | | |
|---|---|---|---|---|
| 97 | BrC | 1 | 2 | low |
| 98 | BrC | 1 | 1 | low |
| 99 | BrC | 1 | 1 | low |
| 100 | BrC | 1 | 1 | low |
| 101 | BrC | 1 | 3 | high |
| 102 | BrC | 0 | 3 | high |
| 103 | BrC | 0 | 3 | low |
| 104 | BrC | 1 | 4 | low |
| 105 | BrC | 1 | 4 | low |
| 106 | BrC | 1 | 4 | low |
| 107 | BrC | 0 | 4 | low |
| 108 | BrC | 0 | 4 | NaN |
| 109 | NSCLC | 1 | 1 | NaN |
| 110 | NSCLC | 1 | 1 | low |
| 111 | NSCLC | 1 | 3 | high |
| 112 | NSCLC | 1 | 3 | high |
| 113 | NSCLC | 0 | 2 | high |
| 114 | NSCLC | 0 | 2 | high |
| 115 | NSCLC | 1 | 2 | high |
| 116 | NSCLC | 1 | 2 | high |
| 117 | NSCLC | 1 | 2 | high |
| 118 | NSCLC | 0 | 1 | high |
| 119 | NSCLC | 0 | 1 | high |
| 120 | NSCLC | 1 | 1 | high |
| 121 | NSCLC | 1 | 3 | high |
| 122 | NSCLC | 0 | 3 | high |
| 123 | NSCLC | 0 | 3 | low |
| 124 | NSCLC | 1 | 4 | low |
| 125 | NSCLC | 1 | 4 | low |
| 126 | NSCLC | 1 | 4 | low |
| 127 | NSCLC | 0 | 4 | high |
| 128 | NSCLC | 0 | 4 | high |
| 129 | NSCLC | 1 | 5 | high |
| 130 | NSCLC | 1 | 5 | low |

| | | | | |
|---|---|---|---|---|
| 131 | OvC | 1 | 1 | low |
| 132 | OvC | 1 | 1 | low |
| 133 | NSCLC | 0 | 4 | high |
| 134 | NSCLC | 0 | 4 | high |
| 135 | NSCLC | 1 | 4 | low |
| 136 | NSCLC | 1 | 3 | low |
| 137 | NSCLC | 1 | 3 | low |
| 138 | NSCLC | 0 | 3 | high |
| 139 | NSCLC | 0 | 4 | high |
| 140 | NSCLC | 1 | 4 | high |
| 141 | OvC | 1 | 4 | low |
| 142 | OvC | 1 | 1 | low |
| 143 | OvC | 1 | 5 | low |
| 144 | OvC | 1 | 5 | high |
| 145 | OvC | 1 | 5 | low |
| 146 | OvC | 1 | 5 | low |
| 147 | OvC | 1 | 5 | low |
| 148 | OvC | 1 | 6 | high |
| 149 | OvC | 1 | 6 | high |
| 150 | OvC | 1 | 6 | high |
| 151 | Tonsil | 0 | 1 | high |
| 152 | Tonsil | 0 | 1 | high |
| 153 | OvC | 1 | 6 | high |
| 154 | OvC | 1 | 6 | low |
| 155 | OvC | 1 | 6 | high |
| 156 | OvC | 0 | 5 | high |
| 157 | OvC | 0 | 5 | low |
| 158 | OvC | 1 | 5 | low |
| 159 | OvC | 1 | 5 | high |
| 160 | OvC | 1 | 5 | high |
| 161 | Tonsil | 0 | 1 | high |
| 162 | Tonsil | 0 | 2 | low |
| 163 | Tonsil | 0 | 2 | low |
| 164 | Tonsil | 0 | 2 | high |

| | | | | |
|---|---|---|---|---|
| 165 | Tonsil | 0 | 3 | high |
| 166 | Tonsil | 0 | 3 | low |
| 167 | Tonsil | 0 | 3 | low |
| 168 | Lymph node | 0 | 1 | high |
| 169 | Lymph node | 0 | 1 | high |
| 170 | Lymph node | 0 | 1 | high |

**Supplementary Table 2:** Normal tissue-microarray metadata.

| Core index | Tissue type |
|---|---|
| 1,2,3 | Pancreas |
| 4,5,6 | Prostate |
| 7,8 | Renal |
| 9,10 | Bladder |
| 11,12 | Ovary |
| 13,14,15 | Lung |
| 16,17,18 | Colon |
| 19,20,21 | Kidney |
| 22,23,24 | Heart |
| 25,26,27 | Tonsil |
| 28,29,30 | Liver |
| 31,32,33 | Spleen |
| 34,35,36 | Lymph node |
| 37,38,39 | Thyroid |
| 40,41,42 | Breast |
| 43,44,45 | Skin |

**Supplementary Table 3:** Gene panel of test iST platforms.

| Xenium breast | Xenium lung | Xenium multi-tissue | MERSCOPE breast | MERSCOPE lung | CosMx 1k |
|---|---|---|---|---|---|
| ABCC11 | ACE | ABCC11 | ABCC11 | ABCC2 | AATK |
| ACTA2 | ACE2 | ACE2 | ADAM9 | ACKR1 | ABL1 |
| ACTG2 | ACKR1 | ACKR1 | ADGRE5 | AGER | ABL2 |
| ADAM9 | ADAM17 | ACTA2 | ADH1B | AGR3 | ACACB |

| | | | | | |
|---|---|---|---|---|---|
| ADGRE5 | ADAM28 | ACTG2 | ADIPOQ | AIF1 | ACE |
| ADH1B | ADAMTS1 | ADAM28 | AGR3 | AKR1B10 | ACKR1 |
| ADIPOQ | ADGRL4 | ADAMTS1 | AIF1 | AKR1C1 | ACKR3 |
| AGR3 | AGER | ADGRE1 | AKR1C1 | AKR1C2 | ACKR4 |
| AIF1 | AGR3 | ADGRL4 | AKR1C3 | ANKRD28 | ACP5 |
| AKR1C1 | AIF1 | ADH1C | ALDH1A3 | ASCL1 | ACTA2 |
| AKR1C3 | ANPEP | ADH4 | ANGPT2 | ATF3 | ACTG2 |
| ALDH1A3 | APOD | ADIPOQ | ANKRD28 | ATF4 | ACVR1 |
| ANGPT2 | APOLD1 | AGER | ANKRD29 | ATF6 | ACVR1B |
| ANKRD28 | AQP9 | AGR3 | ANKRD30A | ATG7 | ACVR2A |
| ANKRD29 | AREG | AHSP | APOBEC3A | ATP2A3 | ACVRL1 |
| ANKRD30A | ARL14 | AIF1 | APOBEC3B | AXL | ADGRA2 |
| APOBEC3A | ASCL1 | ALAS2 | APOC1 | BANK1 | ADGRA3 |
| APOBEC3B | ASCL2 | ALDH1A3 | AQP1 | BAX | ADGRE2 |
| APOC1 | ASCL3 | AMY2A | AR | BCL2 | ADGRE5 |
| AQP1 | ATP1B1 | ANGPT2 | AVPR1A | BCL2L1 | ADGRF1 |
| AQP3 | BAIAP2L1 | ANPEP | BACE2 | BCL2L11 | ADGRF3 |
| AR | BANK1 | APCDD1 | BANK1 | BPIFA1 | ADGRF5 |
| AVPR1A | BCAS1 | APOA5 | BASP1 | C1QC | ADGRG1 |
| BACE2 | BMX | APOBEC3A | C15orf48 | C20orf85 | ADGRG3 |
| BANK1 | CA4 | APOLD1 | C1QA | CCL5 | ADGRG5 |
| BASP1 | CCDC78 | AQP2 | C1QC | CCNA1 | ADGRG6 |
| C15orf48 | CCNA1 | AQP3 | C2orf42 | CCNB2 | ADGRL1 |
| C1QA | CCNB2 | AQP8 | C5orf46 | CCR7 | ADGRL2 |
| C1QC | CCR7 | AQP9 | C6orf132 | CD14 | ADGRL4 |
| C2orf42 | CD14 | AR | CAV1 | CD19 | ADGRV1 |
| C5orf46 | CD163 | ARFGEF3 | CAVIN2 | CD1A | ADIPOQ |
| C6orf132 | CD19 | ASCL1 | CCDC6 | CD1C | ADIRF |
| CAV1 | CD1A | ASCL3 | CCDC80 | CD2 | ADM2 |
| CAVIN2 | CD1C | ASPN | CCL5 | CD247 | AGR2 |
| CCDC6 | CD2 | BAMBI | CCL8 | CD27 | AHI1 |
| CCDC80 | CD24 | BANK1 | CCND1 | CD274 | AHR |
| CCL5 | CD247 | BASP1 | CCPG1 | CD28 | AIF1 |
| CCL8 | CD27 | BBOX1 | CCR7 | CD3D | AKT1 |

| | | | | | |
|---|---|---|---|---|---|
| CCND1 | CD274 | BCL2L11 | CD14 | CD3G | ALCAM |
| CCPG1 | CD28 | BMX | CD163 | CD4 | ALOX5AP |
| CCR7 | CD300E | BTNL9 | CD19 | CD44 | ANGPT1 |
| CD14 | CD34 | C15orf48 | CD247 | CD52 | ANGPT2 |
| CD163 | CD38 | C1orf162 | CD27 | CD68 | ANGPTL1 |
| CD19 | CD3D | C1orf194 | CD274 | CD69 | ANKRD1 |
| CD247 | CD3E | C20orf85 | CD3E | CD79A | ANXA1 |
| CD27 | CD4 | C5orf46 | CD3G | CD79B | ANXA2 |
| CD274 | CD40 | C6orf118 | CD4 | CD86 | ANXA4 |
| CD3E | CD40LG | C7 | CD68 | CD8A | APOA1 |
| CD3G | CD68 | CA4 | CD69 | CD8B | APOC1 |
| CD4 | CD70 | CAPN8 | CD79A | CDH26 | APOD |
| CD68 | CD79A | CAV1 | CD79B | CDK1 | APOE |
| CD69 | CD80 | CAVIN1 | CD80 | CENPF | APP |
| CD79A | CD86 | CAVIN2 | CD83 | CFTR | AQP3 |
| CD79B | CD8A | CCDC39 | CD86 | CGA | AR |
| CD80 | CD8B | CCDC78 | CD8A | CHAC1 | AREG |
| CD83 | CDH1 | CCL19 | CD9 | CHGB | ARF1 |
| CD86 | CDK1 | CCL27 | CD93 | CLDN5 | ARG1 |
| CD8A | CENPF | CCL5 | CDC42EP1 | CPA3 | ARHGDIB |
| CD9 | CFB | CCNB2 | CDH1 | CREB3L4 | ARID5B |
| CD93 | CFTR | CCR2 | CEACAM6 | CRELD2 | ATF3 |
| CDC42EP1 | CHIT1 | CCR7 | CEACAM8 | CTLA4 | ATG10 |
| CDH1 | CLDN5 | CD14 | CENPF | CTNNB1 | ATG12 |
| CEACAM6 | CLEC10A | CD163 | CLEC14A | CXCL13 | ATG5 |
| CEACAM8 | CLEC12A | CD19 | CLEC9A | CXCL14 | ATM |
| CENPF | CLEC4E | CD1A | CLECL1 | CXCL9 | ATP5F1B |
| CLEC14A | CNN1 | CD1C | CLIC6 | CXCR4 | ATP5F1E |
| CLEC9A | COL5A2 | CD1E | CPA3 | CXCR5 | ATR |
| CLECL1 | COL8A1 | CD2 | CRISPLD2 | DCN | AXL |
| CLIC6 | CP | CD247 | CTH | DCTPP1 | AZGP1 |
| CPA3 | CSPG4 | CD27 | CTLA4 | DDIT3 | AZU1 |
| CRISPLD2 | CSTA | CD274 | CTSG | DERL3 | B2M |
| CTH | CTLA4 | CD28 | CTTN | DIRAS3 | B3GNT7 |

| | | | | | |
|---|---|---|---|---|---|
| CTLA4 | CTSL | CD300E | CX3CR1 | DMBT1 | BAG3 |
| CTSG | CTTN | CD34 | CXCL12 | DNAJB9 | BASP1 |
| CTTN | CXCL10 | CD3D | CXCL16 | DUOX1 | BAX |
| CX3CR1 | CXCL13 | CD3E | CXCL5 | EGFR | BBLN |
| CXCL12 | CXCL5 | CD4 | CXCR4 | EHMT1 | BCL2 |
| CXCL16 | CXCL6 | CD5L | CYTIP | EMG1 | BCL2L1 |
| CXCL5 | CXCL9 | CD68 | DAPK3 | EPCAM | BECN1 |
| CXCR4 | CXCR5 | CD69 | DNAAF1 | EREG | BEST1 |
| CYTIP | CXCR6 | CD70 | DNTTIP1 | ERLEC1 | BGN |
| DAPK3 | CYP2F1 | CD79A | DPT | ERN2 | BID |
| DMKN | DAPK2 | CD83 | DSC2 | FAS | BIRC3 |
| DNAAF1 | DCLK1 | CD86 | DSP | FASLG | BIRC5 |
| DNTTIP1 | DES | CD8A | DST | FCER1A | BMP1 |
| DPT | DGKG | CD93 | DUSP2 | FCER1G | BMP2 |
| DSC2 | DIRAS3 | CDH16 | DUSP5 | FCGBP | BMP3 |
| DSP | DMBT1 | CDK1 | EGFL7 | FCGR3A | BMP4 |
| DST | DNAJB9 | CENPF | EGFR | FCN1 | BMP5 |
| DUSP2 | DPP6 | CFAP53 | EIF4EBP1 | FCN3 | BMP7 |
| DUSP5 | DUOX1 | CFB | ENAH | FGFBP2 | BMPR1A |
| EGFL7 | ECSCR | CFHR1 | EPCAM | FKBP11 | BMPR2 |
| EGFR | EGFR | CFHR3 | ERBB2 | FOXP3 | BRAF |
| EIF4EBP1 | EHF | CFTR | ERN1 | GCLM | BRCA1 |
| ENAH | ELF3 | CHGA | ESM1 | GKN2 | BST1 |
| EPCAM | ENAH | CLCA1 | ESR1 | GNG11 | BST2 |
| ERBB2 | EPCAM | CLCA2 | FAM107B | GNLY | BTF3 |
| ERN1 | ETV5 | CLEC10A | FASN | GPR183 | BTG1 |
| ESM1 | F3 | CLEC14A | FBLN1 | GSR | BTK |
| ESR1 | FABP3 | CLEC4E | FCER1A | GZMA | C11orf96 |
| FAM107B | FAM184A | CLECL1 | FCER1G | GZMB | C1QA |
| FAM49A | FAS | CLIC6 | FCGR3A | GZMK | C1QB |
| FASN | FASLG | CNN1 | FGL2 | HAVCR2 | C1QC |
| FBLN1 | FASN | COCH | FLNB | HERPUD1 | C5AR2 |
| FCER1A | FBN1 | COL17A1 | FOXA1 | HLA-DQA1 | CACNA1C |
| FCER1G | FCER1A | COL5A2 | FOXC2 | HLA-DQB1 | CALB1 |

| | | | | | |
|---|---|---|---|---|---|
| FCGR3A | FCGR1A | CPA3 | FOXP3 | HMGA1 | CALD1 |
| FGL2 | FCGR3A | CRHBP | FSTL3 | HMOX1 | CALM1 |
| FLNB | FCMR | CRISPLD2 | GATA3 | HSPA5 | CALM2 |
| FOXA1 | FCN1 | CSF2RA | GJB2 | HYOU1 | CALM3 |
| FOXC2 | FCN3 | CSF3 | GLIPR1 | ICAM1 | CAMP |
| FOXP3 | FGFBP2 | CTLA4 | GNLY | IDH1 | CARMN |
| FSTL3 | FGFR4 | CTSG | GPR183 | IFIT1 | CASP3 |
| GATA3 | FKBP11 | CTSK | GZMA | IFIT2 | CASP8 |
| GJB2 | FOXI1 | CXCL10 | GZMK | IFIT3 | CASR |
| GLIPR1 | FOXJ1 | CXCL2 | HAVCR2 | IL1A | CAV1 |
| GNLY | FOXP3 | CXCL6 | HMGA1 | IL2RA | CCDC80 |
| GPR183 | FSCN1 | CXCL9 | HOXD8 | IL37 | CCL11 |
| GZMA | GJA5 | CXCR4 | HOXD9 | IL4R | CCL13 |
| GZMK | GKN2 | CYP1A1 | HPX | IL7R | CCL15 |
| HAVCR2 | GLCCI1 | CYP2A7 | IGF1 | IRF1 | CCL17 |
| HMGA1 | GLIPR2 | CYP2B6 | IGSF6 | ISG20 | CCL18 |
| HOOK2 | GNG11 | CYP2F1 | IL2RA | ITM2C | CCL19 |
| HOXD8 | GPI | CYP3A4 | IL2RG | KDR | CCL2 |
| HOXD9 | GPR171 | CYP4B1 | IL3RA | KIT | CCL20 |
| HPX | GPR183 | CYTIP | IL7R | KLRB1 | CCL21 |
| IGF1 | GPR34 | DERL3 | ITGAM | KLRC1 | CCL22 |
| IGSF6 | GPX2 | DES | ITGAX | KRT15 | CCL26 |
| IL2RA | GZMA | DIRAS3 | ITM2C | KRT18 | CCL28 |
| IL2RG | GZMB | DMBT1 | JUP | LAG3 | CCL3/L1/L3 |
| IL3RA | GZMK | DNAAF1 | KDR | LEF1 | CCL4/L1/L2 |
| IL7R | HAVCR2 | DNASE1L3 | KIT | LGR5 | CCL5 |
| ITGAM | HIF1A | DPEP1 | KLF5 | LILRA4 | CCL8 |
| ITGAX | HIGD1B | DPT | KLRB1 | LMAN1 | CCND1 |
| ITM2C | HMGCS1 | DST | KLRC1 | LTB | CCR1 |
| JUP | HP | DUSP2 | KLRD1 | LTF | CCR10 |
| KARS | HPGDS | ECSCR | KLRF1 | LUM | CCR2 |
| KDR | ICA1 | EDN1 | KRT23 | MANF | CCR5 |
| KIT | IGF1 | EDNRB | KRT7 | MARCO | CCR7 |
| KLF5 | IGFBP3 | EGFL7 | LAG3 | MCEMP1 | CCRL2 |

| | | | | | |
|---|---|---|---|---|---|
| KLRB1 | IL1RL1 | EGFR | LDHB | MFAP5 | CD14 |
| KLRC1 | IL7R | EHF | LGALSL | MGST1 | CD163 |
| KLRD1 | IQGAP2 | ELF5 | LILRA4 | MKI67 | CD164 |
| KLRF1 | IRF8 | EPCAM | LPL | MMP10 | CD19 |
| KRT14 | ITGAM | ERBB2 | LPXN | MMP12 | CD1C |
| KRT23 | ITGB4 | ERG | LRRC15 | MS4A1 | CD2 |
| KRT5 | KCNK3 | ESR1 | LTB | MS4A7 | CD209 |
| KRT6B | KDR | FAS | LUM | MUC5B | CD22 |
| KRT7 | KIT | FBLN1 | LY86 | MYC | CD24 |
| KRT8 | KLF5 | FBN1 | LYPD3 | MYDGF | CD27 |
| LAG3 | KLK11 | FCER1A | LYZ | NAPSA | CD274 |
| LARS | KLRB1 | FCGR1A | MAP3K8 | NFKB1 | CD276 |
| LDHB | KLRC1 | FCGR3A | MDM2 | NHSL2 | CD28 |
| LGALSL | KLRD1 | FCN1 | MKI67 | NKG7 | CD300A |
| LILRA4 | KRT15 | FCN2 | MLPH | NKX2-1 | CD33 |
| LPL | KRT7 | FGFBP1 | MMP1 | NUCB2 | CD34 |
| LPXN | LAG3 | FGFBP2 | MMP12 | NUTF2 | CD36 |
| LRRC15 | LAMC3 | FGL2 | MMP2 | OAS2 | CD37 |
| LTB | LCK | FHL2 | MMRN2 | OAS3 | CD38 |
| LUM | LGALS3BP | FKBP11 | MNDA | PAEP | CD3D |
| LY86 | LGR5 | FOXA1 | MRC1 | PCNA | CD3E |
| LYPD3 | LGR6 | FOXI1 | MS4A1 | PDCD1 | CD3G |
| LYZ | LILRA4 | FOXJ1 | MUC6 | PDIA3 | CD4 |
| MAP3K8 | LILRA5 | FOXP3 | MYBPC1 | PDIA4 | CD40 |
| MDM2 | LILRB2 | FSTL3 | MYLK | PDIA6 | CD40LG |
| MKI67 | LILRB4 | FXYD2 | MYO5B | PECAM1 | CD44 |
| MLPH | LMOD1 | GATA2 | MZB1 | PGC | CD47 |
| MMP1 | LTBP2 | GATM | NCAM1 | PIM2 | CD48 |
| MMP12 | LTF | GCG | NDUFA4L2 | PLPP5 | CD52 |
| MMP2 | LYVE1 | GDF15 | NKG7 | POSTN | CD53 |
| MMRN2 | MALL | GEM | NOSTRIN | PRDX4 | CD55 |
| MNDA | MAP7 | GHRL | OCIAD2 | PTPRC | CD58 |
| MRC1 | MARCO | GKN2 | OPRPN | RACGAP1 | CD59 |
| MS4A1 | MCEMP1 | GLIPR1 | OXTR | RAMP2 | CD5L |

| | | | | | |
|---|---|---|---|---|---|
| MUC6 | MEDAG | GLYATL1 | PCLAF | RHOA | CD63 |
| MYBPC1 | MET | GNG11 | PDCD1 | RNASE1 | CD68 |
| MYH11 | MFAP5 | GNLY | PDCD1LG2 | S100A12 | CD69 |
| MYLK | MIS18BP1 | GPC1 | PDE4A | SAA2 | CD70 |
| MYO5B | MKI67 | GPC3 | PDGFRA | SCG2 | CD74 |
| MZB1 | MMP12 | GPR183 | PDGFRB | SEC11C | CD79A |
| NARS | MMP9 | GPRC5A | PDK4 | SELENOS | CD80 |
| NCAM1 | MMRN1 | GPX2 | PECAM1 | SFRP2 | CD81 |
| NDUFA4L2 | MPEG1 | GYPA | PELI1 | SFTA2 | CD83 |
| NKG7 | MS4A1 | GYPB | PGR | SFTPD | CD84 |
| NOSTRIN | MS4A2 | GZMA | PIM1 | SLC1A3 | CD86 |
| OCIAD2 | MS4A4A | GZMB | PLD4 | SLC25A37 | CD8A |
| OPRPN | MTUS1 | GZMK | POSTN | SLC25A4 | CD8B |
| OXTR | MUC1 | HAMP | PPARG | SLC2A1 | CD9 |
| PCLAF | MUC5B | HAVCR2 | PRDM1 | SLC7A11 | CD93 |
| PDCD1 | MYC | HEMGN | PRF1 | SMAD4 | CDH1 |
| PDCD1LG2 | MYH11 | HEPACAM2 | PTN | SNAI1 | CDH11 |
| PDE4A | MYO6 | HES4 | PTPRC | SNAI2 | CDH19 |
| PDGFRA | MZB1 | HIGD1B | PTRHD1 | SNCA | CDH5 |
| PDGFRB | NCEH1 | HLA-DQB2 | RAB30 | SOD2 | CDKN1A |
| PDK4 | NFKB1 | HMGCS2 | RAMP2 | SOX9 | CDKN3 |
| PECAM1 | NID1 | HPGDS | RAPGEF3 | SPARCL1 | CEACAM1 |
| PELI1 | NKG7 | HPX | RHOH | SPCS2 | CEACAM6 |
| PGR | NTN4 | IGF1 | RORC | SPCS3 | CELSR1 |
| PIGR | NTRK2 | IGSF6 | RTKN2 | SSR3 | CELSR2 |
| PIM1 | OTUD7B | IL1R2 | RUNX1 | STAT1 | CENPF |
| PLD4 | P2RX1 | IL1RL1 | SCD | STAT6 | CFD |
| POSTN | PAMR1 | IL2RA | SCGB2A1 | TCL1A | CFLAR |
| PPARG | PCNA | IL3RA | SDC4 | TGFB1 | CHEK1 |
| PRDM1 | PCOLCE2 | IL7R | SEC11C | TNFRSF13C | CHEK2 |
| PRF1 | PDCD1 | INMT | SEC24A | TNFRSF17 | CHI3L1 |
| PTGDS | PDCD1LG2 | INS | SELL | TNFRSF9 | CIDEA |
| PTN | PDGFRA | IRF8 | SERPINB9 | TOP1 | CIITA |
| PTPRC | PDGFRB | KCNK3 | SFRP1 | TOP2A | CLCF1 |

| | | | | | |
|---|---|---|---|---|---|
| PTRHD1 | PDPN | KCNMA1 | SFRP4 | TP63 | CLDN4 |
| QARS | PEBP4 | KIT | SH3YL1 | TRAC | CLEC10A |
| RAB30 | PI3 | KLK11 | SLAMF1 | TTC19 | CLEC12A |
| RAMP2 | PIM1 | KLRB1 | SLAMF7 | UBE2J1 | CLEC14A |
| RAPGEF3 | PIM2 | KLRC1 | SLC25A37 | UBE2S | CLEC1A |
| RHOH | PLA2G2A | KLRD1 | SLC5A6 | UCHL3 | CLEC2B |
| RORC | PLA2G4F | KNG1 | SMAP2 | UGDH | CLEC2D |
| RTKN2 | PLA2G7 | KRT20 | SMS | UQCRHL | CLEC4A |
| RUNX1 | PLN | KRT7 | SNAI1 | VEGFA | CLEC4D |
| S100A14 | PLVAP | LAG3 | SOX17 | VPREB3 | CLEC4E |
| S100A4 | POU2AF1 | LAMP3 | SOX18 | WFDC2 | CLEC5A |
| S100A8 | PROX1 | LGI4 | SPIB | ZEB1 | CLEC7A |
| SCD | PTGS1 | LGR5 | SQLE | | CLOCK |
| SCGB2A1 | RAMP2 | LIF | SRPK1 | | CLU |
| SDC4 | RARRES1 | LILRA4 | SSTR2 | | CMKLR1 |
| SEC11C | RBP4 | LILRA5 | STC1 | | CNTFR |
| SEC24A | RERGL | LILRB2 | SVIL | | COL11A1 |
| SELL | RETN | LILRB4 | TAC1 | | COL12A1 |
| SERHL2 | RGS5 | LPL | TACSTD2 | | COL14A1 |
| SERPINA3 | RND1 | LTBP2 | TCEAL7 | | COL15A1 |
| SERPINB9 | RUNX3 | LY6D | TCF4 | | COL16A1 |
| SFRP1 | S100A12 | LY86 | TCF7 | | COL17A1 |
| SFRP4 | S100A7 | LYVE1 | TCIM | | COL18A1 |
| SH3YL1 | S100B | MALL | TCL1A | | COL1A1 |
| SLAMF1 | SAMD3 | MAMDC2 | TENT5C | | COL1A2 |
| SLAMF7 | SCEL | MARCO | TFAP2A | | COL21A1 |
| SLC25A37 | SEC11C | MCEMP1 | THAP2 | | COL27A1 |
| SLC5A6 | SELE | MCF2L | TIFA | | COL3A1 |
| SMAP2 | SELL | MDM2 | TIGIT | | COL4A1 |
| SMS | SELP | MEDAG | TIMP4 | | COL4A2 |
| SNAI1 | SEMA3B | MEF2C | TMEM147 | | COL4A5 |
| SOX17 | SEMA3C | MEST | TNFRSF17 | | COL5A1 |
| SOX18 | SERPINA3 | MET | TOMM7 | | COL5A2 |
| SPIB | SFRP2 | MFAP5 | TOP2A | | COL5A3 |

| | | | | |
|---|---|---|---|---|
| SQLE | SFTA2 | MKI67 | TPD52 | COL6A1 |
| SRPK1 | SFTPD | MLANA | TRAC | COL6A2 |
| SSTR2 | SHANK3 | MLPH | TRAPPC3 | COL6A3 |
| STC1 | SLC15A2 | MMRN1 | TRIB1 | COL8A1 |
| SVIL | SLC18A2 | MMRN2 | TUBA4A | COL9A2 |
| TAC1 | SLC1A3 | MNDA | TUBB2B | COL9A3 |
| TACSTD2 | SLC2A1 | MPEG1 | UCP1 | COTL1 |
| TCEAL7 | SLC7A11 | MRC1 | USP53 | COX4I2 |
| TCF4 | SLIT3 | MS4A1 | VOPP1 | CPA3 |
| TCF7 | SMIM24 | MS4A2 | VWF | CPB1 |
| TCIM | SOX2 | MS4A4A | ZEB1 | CRIP1 |
| TCL1A | SOX9 | MS4A6A | ZEB2 | CRP |
| TENT5C | SPDEF | MTRNR2L11 | ZNF562 | CRYAB |
| TFAP2A | SPIB | MYBPC1 | | CSF1 |
| THAP2 | STAT4 | MYC | | CSF1R |
| TIFA | STC1 | MYH11 | | CSF2 |
| TIGIT | STEAP4 | MYLK | | CSF2RA |
| TIMP4 | SVEP1 | MZB1 | | CSF2RB |
| TMEM147 | SYK | NAT8 | | CSF3 |
| TNFRSF17 | TACSTD2 | NKG7 | | CSF3R |
| TOMM7 | TC2N | NPDC1 | | CSHL1 |
| TOP2A | TCL1A | NTN4 | | CSK |
| TPD52 | THBS2 | OGN | | CSPG4 |
| TPSAB1 | THY1 | OPRPN | | CST7 |
| TRAC | TM4SF18 | PCNA | | CSTB |
| TRAF4 | TM4SF4 | PCOLCE | | CTLA4 |
| TRAPPC3 | TMC5 | PCP4 | | CTNNB1 |
| TRIB1 | TMEM100 | PCSK2 | | CTSD |
| TUBA4A | TMPRSS2 | PDCD1 | | CTSG |
| TUBB2B | TNFRSF13B | PDGFRA | | CTSW |
| UCP1 | TNFRSF13C | PDGFRB | | CUZD1 |
| USP53 | TNFRSF17 | PDPN | | CX3CL1 |
| VOPP1 | TNFRSF18 | PEBP4 | | CX3CR1 |
| VWF | TOP2A | PECAM1 | | CXCL1/2/3 |

| | | | |
|---|---|---|---|
| WARS | TP63 | PGR | CXCL10 |
| ZEB1 | TP73 | PLA2G7 | CXCL12 |
| ZEB2 | TREM2 | PLAC9 | CXCL13 |
| ZNF562 | TRPC6 | PLCG2 | CXCL14 |
| | TSPAN8 | PLD4 | CXCL16 |
| | UBE2C | PLIN4 | CXCL17 |
| | UPK1B | PMP22 | CXCL5 |
| | UPK3B | PPARG | CXCL8 |
| | VSIG4 | PPP1R1A | CXCL9 |
| | VWF | PPP1R1B | CXCR1 |
| | WFS1 | PPY | CXCR2 |
| | WNT2 | PRDM1 | CXCR3 |
| | WT1 | PRF1 | CXCR4 |
| | | PRG4 | CXCR5 |
| | | PROX1 | CXCR6 |
| | | PTGDS | CYP1B1 |
| | | PTN | CYP2U1 |
| | | PTPRC | CYSTM1 |
| | | PVALB | CYTOR |
| | | RAMP2 | DCN |
| | | RAPGEF3 | DDC |
| | | RBP5 | DDIT3 |
| | | RERGL | DDR1 |
| | | RETN | DDR2 |
| | | RGS16 | DDX58 |
| | | RIDA | DHRS2 |
| | | RND1 | DLL1 |
| | | RTKN2 | DLL4 |
| | | S100A1 | DMBT1 |
| | | S100A12 | DNMT1 |
| | | SCGB2A1 | DNMT3A |
| | | SCGN | DPP4 |
| | | SELE | DPT |
| | | SELL | DST |

| | |
|---|---|
| SEMA3C | DUSP1 |
| SERPINB2 | DUSP2 |
| SERPINB3 | DUSP4 |
| SERPINB9 | DUSP5 |
| SFRP2 | DUSP6 |
| SFRP4 | EFNA1 |
| SFTA2 | EFNA4 |
| SH2D3C | EFNA5 |
| SLAMF1 | EFNB1 |
| SLAMF7 | EFNB2 |
| SLC18A2 | EGF |
| SLC22A8 | EGFR |
| SLC26A2 | EIF5A/L1 |
| SLC26A3 | ELANE |
| SLC4A1 | EMP3 |
| SMIM24 | ENG |
| SMYD2 | ENO1 |
| SNAI1 | ENTPD1 |
| SNCA | EOMES |
| SNCG | EPCAM |
| SNTN | EPHA2 |
| SOX17 | EPHA3 |
| SOX18 | EPHA4 |
| SOX2 | EPHA7 |
| SPDEF | EPHB2 |
| SPI1 | EPHB3 |
| SPIB | EPHB4 |
| SRPX | EPHB6 |
| SST | EPOR |
| STC1 | ERBB2 |
| STC2 | ERBB3 |
| STEAP4 | ESAM |
| TAC1 | ESR1 |
| TAT | ETS1 |

| | |
|---|---|
| TBX3 | ETV4 |
| TCF15 | ETV5 |
| TCF4 | EZH2 |
| TCIM | EZR |
| TCL1A | FABP4 |
| TENT5C | FABP5 |
| TFF2 | FAM30A |
| TFPI | FAP |
| THAP2 | FAS |
| THBS2 | FASLG |
| THY1 | FASN |
| TIMP4 | FAU |
| TM4SF18 | FCER1G |
| TM4SF4 | FCGBP |
| TMC5 | FCGR3A/B |
| TMEM100 | FCRLA |
| TMEM174 | FES |
| TMEM52B | FFAR2 |
| TNC | FFAR3 |
| TNFRSF13B | FFAR4 |
| TNFRSF17 | FGF1 |
| TNFRSF9 | FGF12 |
| TOP2A | FGF13 |
| TRAC | FGF18 |
| TREM2 | FGF2 |
| TSPAN19 | FGF7 |
| UBE2C | FGF9 |
| UMOD | FGFR1 |
| UPK3B | FGFR2 |
| VCAN | FGFR3 |
| VSIG4 | FGG |
| VWA5A | FGR |
| VWF | FHIT |
| | FKBP11 |

| |
|---|
| FKBP5 |
| FLT1 |
| FLT3LG |
| FN1 |
| FOS |
| FOXF1 |
| FOXP3 |
| FPR1 |
| FYB1 |
| FYN |
| FZD1 |
| FZD3 |
| FZD4 |
| FZD5 |
| FZD6 |
| FZD7 |
| FZD8 |
| G0S2 |
| G6PD |
| GADD45B |
| GAS6 |
| GATA3 |
| GC |
| GCG |
| GDF15 |
| GLUD1 |
| GLUL |
| GNLY |
| GPBAR1 |
| GPER1 |
| GPNMB |
| GPR183 |
| GPX1 |
| GPX3 |

| |
|---|
| GSK3B |
| GSN |
| GSTP1 |
| GZMA |
| GZMB |
| GZMH |
| GZMK |
| H2AZ1 |
| H4C3 |
| HAVCR2 |
| HBA1/2 |
| HBB |
| HCAR2/3 |
| HCK |
| HCST |
| HDAC1 |
| HDAC11 |
| HDAC3 |
| HDAC4 |
| HDAC5 |
| HEXB |
| HEY1 |
| HGF |
| HIF1A |
| HILPDA |
| HLA-DPA1 |
| HLA-DPB1 |
| HLA-DQA1 |
| HLA-DQB1/2 |
| HLA-DRA |
| HLA-DRB |
| HMGB2 |
| HMGCS1 |
| HMGN2 |

| |
|---|
| HPGDS |
| HSD17B2 |
| HSP90AA1 |
| HSP90AB1 |
| HSP90B1 |
| HSPA1A/B |
| HSPB1 |
| HSPG2 |
| HTT |
| IAPP |
| ICA1 |
| ICAM1 |
| ICAM2 |
| ICAM3 |
| ICOS |
| ICOSLG |
| IDO1 |
| IER3 |
| IFI27 |
| IFI44L |
| IFI6 |
| IFIH1 |
| IFIT1 |
| IFIT3 |
| IFITM1 |
| IFITM3 |
| IFNA1/13 |
| IFNAR1 |
| IFNAR2 |
| IFNG |
| IFNGR1 |
| IFNGR2 |
| IFNL2/3 |
| IGF1 |

| |
|---|
| IGF1R |
| IGF2 |
| IGF2R |
| IGFBP3 |
| IGFBP5 |
| IGFBP6 |
| IGFBP7 |
| IGHA1 |
| IGHD |
| IGHG1 |
| IGHG2 |
| IGHM |
| IGKC |
| IKZF3 |
| IL10 |
| IL10RA |
| IL10RB |
| IL11 |
| IL11RA |
| IL12A |
| IL12B |
| IL12RB1 |
| IL12RB2 |
| IL13RA1 |
| IL15 |
| IL15RA |
| IL16 |
| IL17A |
| IL17B |
| IL17D |
| IL17RA |
| IL17RB |
| IL17RE |
| IL18 |

| |
|---|
| IL18R1 |
| IL1A |
| IL1B |
| IL1R1 |
| IL1R2 |
| IL1RAP |
| IL1RL1 |
| IL1RN |
| IL2 |
| IL20 |
| IL20RA |
| IL22RA1 |
| IL23A |
| IL24 |
| IL27RA |
| IL2RA |
| IL2RB |
| IL2RG |
| IL32 |
| IL33 |
| IL34 |
| IL36G |
| IL3RA |
| IL4R |
| IL6 |
| IL6R |
| IL6ST |
| IL7 |
| IL7R |
| INHA |
| INHBA |
| INHBB |
| INS |
| INSIG1 |

| |
|---|
| INSR |
| IRF3 |
| IRF4 |
| ISG15 |
| ITGA1 |
| ITGA2 |
| ITGA3 |
| ITGA5 |
| ITGA6 |
| ITGA8 |
| ITGA9 |
| ITGAE |
| ITGAL |
| ITGAM |
| ITGAV |
| ITGAX |
| ITGB1 |
| ITGB2 |
| ITGB4 |
| ITGB5 |
| ITGB6 |
| ITGB8 |
| ITK |
| ITM2A |
| ITM2B |
| JAG1 |
| JAK1 |
| JAK2 |
| JCHAIN |
| JUN |
| JUNB |
| KDR |
| KIT |
| KITLG |

| |
|---|
| KLF2 |
| KLK3 |
| KLRB1 |
| KLRF1 |
| KLRK1 |
| KRAS |
| KRT1 |
| KRT10 |
| KRT13 |
| KRT14 |
| KRT15 |
| KRT16 |
| KRT17 |
| KRT18 |
| KRT19 |
| KRT20 |
| KRT23 |
| KRT4 |
| KRT5 |
| KRT6A/B/C |
| KRT7 |
| KRT8 |
| KRT80 |
| KRT86 |
| LAG3 |
| LAIR1 |
| LAMA4 |
| LAMP2 |
| LAMP3 |
| LCN2 |
| LDB2 |
| LDHA |
| LDLR |
| LEFTY1 |

| |
|---|
| LEP |
| LGALS1 |
| LGALS3 |
| LGALS3BP |
| LGALS9 |
| LGR5 |
| LIF |
| LIFR |
| LINC01781 |
| LINC01857 |
| LINC02446 |
| LMNA |
| LPAR5 |
| LTB |
| LTBR |
| LTF |
| LUM |
| LY6D |
| LY75 |
| LYN |
| LYVE1 |
| LYZ |
| MAF |
| MALAT1 |
| MAML2 |
| MAP1LC3B/2 |
| MAP2K1 |
| MAPK13 |
| MAPK14 |
| MARCKSL1 |
| MARCO |
| MB |
| MECOM |
| MEG3 |

| |
|---|
| MERTK |
| MET |
| MFAP5 |
| MGP |
| MHC I |
| MIF |
| MIR4435-2HG |
| MKI67 |
| MMP1 |
| MMP12 |
| MMP14 |
| MMP19 |
| MMP2 |
| MMP7 |
| MMP9 |
| MPO |
| MRC1 |
| MRC2 |
| MS4A1 |
| MS4A4A |
| MS4A6A |
| MSMB |
| MSR1 |
| MST1R |
| MT1X |
| MT2A |
| MTOR |
| MX1 |
| MXRA8 |
| MYC |
| MYH11 |
| MYH6 |
| MYL12A |
| MYL4 |

| |
|---|
| MYL7 |
| MYL9 |
| MZB1 |
| MZT2A/B |
| NACA |
| NANOG |
| NCAM1 |
| NCR1 |
| NDRG1 |
| NDUFA4L2 |
| NEAT1 |
| NELL2 |
| NFKB1 |
| NFKBIA |
| NGFR |
| NKG7 |
| NLRC4 |
| NLRC5 |
| NLRP1 |
| NLRP2 |
| NLRP3 |
| NOD2 |
| NOSIP |
| NOTCH1 |
| NOTCH2 |
| NOTCH3 |
| NPPC |
| NPR1 |
| NPR2 |
| NPR3 |
| NR1H2 |
| NR1H3 |
| NR2F2 |
| NR3C1 |

| |
|---|
| NRG1 |
| NRXN1 |
| NRXN3 |
| NTRK2 |
| NUPR1 |
| NUSAP1 |
| OAS1 |
| OAS2 |
| OAS3 |
| OASL |
| OLFM4 |
| OLR1 |
| OSM |
| OSMR |
| P2RX5 |
| PARP1 |
| PCNA |
| PDCD1 |
| PDCD1LG2 |
| PDGFA |
| PDGFB |
| PDGFC |
| PDGFD |
| PDGFRA |
| PDGFRB |
| PDS5A |
| PECAM1 |
| PF4/V1 |
| PFN1 |
| PGF |
| PGK1 |
| PGR |
| PHLDA2 |
| PIGR |

| |
|---|
| PLAC8 |
| PLAC9 |
| PLCG1 |
| PLD3 |
| PNOC |
| POU5F1 |
| PPARA |
| PPARD |
| PPARG |
| PPIA |
| PRF1 |
| PROX1 |
| PRSS2 |
| PRTN3 |
| PSAP |
| PSCA |
| PSD3 |
| PTEN |
| PTGDR2 |
| PTGDS |
| PTGES |
| PTGES2 |
| PTGES3 |
| PTGIS |
| PTGS1 |
| PTGS2 |
| PTK2 |
| PTK6 |
| PTPRC |
| PTPRCAP |
| PTTG1 |
| PXDN |
| QRFPR |
| RAC1 |

| |
|---|
| RAC2 |
| RACK1 |
| RAG1 |
| RAMP1 |
| RAMP2 |
| RAMP3 |
| RARA |
| RARB |
| RARG |
| RARRES1 |
| RARRES2 |
| RB1 |
| RBM47 |
| RBPJ |
| REG1A |
| RELA |
| RELT |
| RGCC |
| RGS1 |
| RGS13 |
| RGS2 |
| RGS5 |
| RNF43 |
| ROR1 |
| RORA |
| RPL21 |
| RPL22 |
| RPL32 |
| RPL34 |
| RPL37 |
| RPS4Y1 |
| RSPO3 |
| RUNX3 |
| RXRA |

| |
|---|
| RXRB |
| RYK |
| RYR2 |
| S100A10 |
| S100A2 |
| S100A4 |
| S100A6 |
| S100A8 |
| S100A9 |
| S100B |
| S100P |
| SAA1/2 |
| SAT1 |
| SCG5 |
| SCGB3A1 |
| SEC23A |
| SEC61G |
| SELENOP |
| SELL |
| SELPLG |
| SERPINA1 |
| SERPINA3 |
| SERPINB5 |
| SERPINH1 |
| SFN |
| SH3BGRL3 |
| SIGIRR |
| SLA |
| SLC2A1 |
| SLC40A1 |
| SLCO2B1 |
| SLPI |
| SMAD2 |
| SMAD3 |

| |
|---|
| SMAD4 |
| SMARCB1 |
| SMO |
| SNAI1 |
| SNAI2 |
| SOD1 |
| SOD2 |
| SORBS1 |
| SOSTDC1 |
| SOX2 |
| SOX4 |
| SOX9 |
| SPARCL1 |
| SPINK1 |
| SPOCK2 |
| SPP1 |
| SPRY2 |
| SPRY4 |
| SQLE |
| SQSTM1 |
| SRC |
| SREBF1 |
| SRGN |
| SRSF2 |
| SST |
| ST6GAL1 |
| ST6GALNAC3 |
| STAT1 |
| STAT3 |
| STAT4 |
| STAT5A |
| STAT5B |
| STAT6 |
| STMN1 |

| |
| --- |
| SYK |
| TACSTD2 |
| TAGLN |
| TAP1 |
| TAP2 |
| TBX21 |
| TCAP |
| TCF7 |
| TCL1A |
| TEK |
| TFEB |
| TGFB1 |
| TGFB2 |
| TGFB3 |
| TGFBI |
| TGFBR1 |
| TGFBR2 |
| THBS1 |
| THBS2 |
| THSD4 |
| TIE1 |
| TIGIT |
| TIMP1 |
| TLR1 |
| TLR2 |
| TLR3 |
| TLR4 |
| TLR5 |
| TLR7 |
| TLR8 |
| TM4SF1 |
| TNF |
| TNFAIP6 |
| TNFRSF10A |

| |
|---|
| TNFRSF10B |
| TNFRSF10D |
| TNFRSF11A |
| TNFRSF11B |
| TNFRSF12A |
| TNFRSF13B |
| TNFRSF14 |
| TNFRSF17 |
| TNFRSF18 |
| TNFRSF19 |
| TNFRSF1A |
| TNFRSF1B |
| TNFRSF21 |
| TNFRSF4 |
| TNFRSF9 |
| TNFSF10 |
| TNFSF12 |
| TNFSF13B |
| TNFSF14 |
| TNFSF15 |
| TNFSF4 |
| TNFSF8 |
| TNFSF9 |
| TNNC1 |
| TNNT2 |
| TNXA/B |
| TOP2A |
| TOX |
| TP53 |
| TPI1 |
| TPM1 |
| TPM2 |
| TPSAB1/B2 |
| TPT1 |

| |
|---|
| TSC22D1 |
| TSHZ2 |
| TTN |
| TTR |
| TUBB |
| TUBB4B |
| TWIST1 |
| TWIST2 |
| TXK |
| TYK2 |
| TYMS |
| TYROBP |
| UBA52 |
| UBE2C |
| UPK3A |
| VCAM1 |
| VCAN |
| VEGFA |
| VEGFB |
| VEGFC |
| VEGFD |
| VHL |
| VIM |
| VPREB3 |
| VSIR |
| VTN |
| VWA1 |
| VWF |
| WIF1 |
| WNT10B |
| WNT11 |
| WNT3 |
| WNT5A |
| WNT5B |

| |
|---|
| WNT7A |
| WNT7B |
| WNT9A |
| XBP1 |
| XCL1/2 |
| XKR4 |
| YBX3 |
| YES1 |
| ZBTB16 |
| ZFP36 |

**Supplementary Table 4:** Metrics by each run, combination of platform, panel, and TMA.

| TMA | Platform | Panel | Data collection date (Year: 2023) | Days after slicing | Cores with data | Cores with data (%) | Cell count (k) | Total transcripts (million) | Transcript per cell | Cells per 1000µm^2 | Good cells (%, transcripts>10 per cell) | Good cells (%, transcripts>20 per cell) |
|---|---|---|---|---|---|---|---|---|---|---|---|---|
| Tumor | Xenium | Breast | May-10 | 6 | 169 | 97.7 | 399.3 | 46.6 | 116.6 | 7.4 | 97.4 | 92.8 |
| | Xenium | Multi-tissue | May-25 | 21 | 159 | 91.9 | 379.5 | 33.8 | 89.2 | 7.2 | 97.1 | 91.7 |
| | Xenium | Lung | Jun-16 | 43 | 149 | 86.1 | 360.8 | 28.2 | 78.2 | 7.1 | 95.1 | 86.7 |
| | CosMx | 1K | Jun-17 | 44 | 163 | 94.2 | 240.2 | 22.3 | 92.8 | 5.0 | 93.2 | 83.4 |
| | MERSCOPE | Breast | Jul-17 | 74 | 107 | 61.8 | 232.4 | 13.2 | 56.8 | 6.1 | 68.5 | 57.9 |
| | MERSCOPE | Lung | May-25 | 21 | 96 | 55.5 | 274.0 | 2.4 | 8.8 | 4.8 | 25.8 | 12.3 |
| Normal | Xenium | Breast | May-10 | 6 | 48 | 100.0 | 300.9 | 14.7 | 49.0 | 2.2 | 84.0 | 71.5 |
| | Xenium | Multi-tissue | May-25 | 21 | 48 | 100.0 | 292.7 | 11.7 | 39.9 | 2.3 | 84.3 | 68.5 |
| | Xenium | Lung | Jun-16 | 43 | 48 | 100.0 | 252.6 | 8.5 | 33.5 | 2.0 | 77.4 | 58.8 |
| | CosMx | 1K | Jun-17 | 44 | 47 | 97.9 | 178.4 | 8.4 | 47.0 | 2.2 | 86.8 | 67.1 |
| | MERSCOPE | Breast | Jul-17 | 74 | 38 | 79.2 | 164.0 | 0.2 | 1.5 | 0.2 | 2.4 | 0.6 |
| | MERSCOPE | Lung | May-23 | 19 | 45 | 93.8 | 255.3 | 0.1 | 0.4 | 0.3 | 0.1 | 0.0 |

**Supplementary Table 5:** Number of genes detected two standard deviations above the average expression of the negative control probes. The max value of each tissue type was bolded.

| Platform (panel) Tissue Type\ | CosMx 1k | MERSCOPE breast | MERSCOPE lung | Xenium breast | Xenium lung | Xenium multi-tissue |
|---|---|---|---|---|---|---|
| Bladder | **246** | 101 | 123 | 154 | 243 | 208 |
| BrC | **526** | 138 | 90 | 265 | 271 | 346 |
| Breast | **431** | 98 | 91 | 230 | 270 | 347 |
| CRC | **503** | 154 | 122 | 252 | 274 | 343 |
| Colon | 241 | 101 | 74 | 210 | 267 | **311** |
| HNSCC | **394** | 130 | 107 | 257 | 235 | 332 |
| Heart | 218 | 29 | 46 | 162 | 185 | **243** |
| Kidney | 165 | 155 | 118 | 272 | 276 | **359** |
| Liver | 192 | 112 | 91 | 246 | 272 | **346** |
| Lung | 170 | 27 | 41 | 228 | 276 | **325** |
| Lymph node | 197 | 85 | 53 | 190 | 274 | **336** |
| Mel | **412** | 106 | 77 | 265 | 265 | 343 |
| NSCLC | **406** | 145 | 95 | 267 | 281 | 345 |
| OvC | **503** | 180 | 109 | 262 | 279 | 357 |
| Ovary | **566** | 203 | 144 | 270 | 286 | 347 |
| Prostate | **505** | 185 | 150 | 247 | 287 | 356 |
| Skin | **417** | 66 | 95 | 270 | 281 | 367 |
| Spleen | 157 | 21 | 48 | 279 | 283 | **369** |
| Thyroid | **675** | 136 | 129 | 275 | 287 | 369 |
| Tonsil | **569** | 144 | 126 | 246 | 261 | 361 |

**Supplementary Table 6:** Number of segmented cells per 1000 μm$^2$ by tissue type and by platform x panel combination.

| Platform (panel) Tissue Type\ | CosMx 1k | MERSCOPE breast | MERSCOPE lung | Xenium breast | Xenium lung | Xenium multi-tissue |
|---|---|---|---|---|---|---|
| Bladder | 1.83 | 0.70 | 0.36 | 1.53 | 1.17 | 1.65 |
| BrC | 4.65 | 6.87 | 4.81 | 7.48 | 7.33 | 7.73 |
| Breast | 1.91 | 0.06 | 0.18 | 0.46 | 1.00 | 0.65 |
| CRC | 5.26 | 6.49 | 5.34 | 8.62 | 7.69 | 8.41 |
| Colon | 2.40 | 0.43 | 0.27 | 3.29 | 2.64 | 3.21 |
| HNSCC | 4.44 | 4.61 | 2.70 | 6.43 | 5.53 | 4.76 |
| Heart | 0.71 | 0.20 | 0.20 | 1.04 | 0.94 | 0.95 |
| Kidney | 2.95 | 1.27 | 0.40 | 4.00 | 3.42 | 3.83 |
| Liver | 2.23 | 0.57 | 0.85 | 2.14 | 1.96 | 2.06 |
| Lung | 0.29 | 0.07 | 0.03 | 0.70 | 0.54 | 0.62 |
| Lymph node | 0.51 | 0.08 | 0.06 | 0.19 | 0.19 | 0.23 |
| Mel | 7.27 | 5.02 | 2.77 | 9.66 | 9.01 | 7.78 |
| NSCLC | 4.56 | 4.83 | 3.78 | 6.70 | 6.15 | 7.54 |
| OvC | 6.43 | 11.40 | 9.69 | 13.29 | 12.38 | 12.07 |
| Ovary | 2.49 | 1.20 | 0.80 | 2.74 | 2.19 | 2.52 |
| Prostate | 3.32 | 0.79 | 0.89 | 3.68 | 3.81 | 3.98 |
| Skin | 2.18 | 0.12 | 0.13 | 1.73 | 1.36 | 1.61 |
| Spleen | 9.57 | 1.69 | 1.63 | 15.39 | 10.98 | 15.66 |
| Thyroid | 0.71 | 0.13 | 0.53 | 2.68 | 2.91 | 2.34 |
| Tonsil | 10.89 | 0.45 | 2.79 | 17.39 | 13.87 | 15.27 |

**Supplementary Table 7:** Area of segmented cells ($\mu m^2$) by tissue type and by platform x panel combination.

| Platform (panel) Tissue Type\ | CosMx 1k | MERSCOPE breast | MERSCOPE lung | Xenium breast | Xenium lung | Xenium multi-tissue |
|---|---|---|---|---|---|---|
| Bladder | 190.70 | 85.59 | 79.70 | 343.30 | 345.11 | 342.01 |
| BrC | 125.11 | 72.00 | 70.84 | 112.78 | 112.78 | 102.25 |
| Breast | 143.24 | 75.28 | 72.78 | 169.79 | 130.16 | 151.72 |
| CRC | 112.28 | 63.51 | 59.65 | 95.82 | 94.38 | 93.45 |
| Colon | 149.24 | 77.97 | 76.65 | 148.86 | 161.37 | 158.59 |
| HNSCC | 147.56 | 86.03 | 79.98 | 153.17 | 162.52 | 152.52 |
| Heart | 220.02 | 91.58 | 71.24 | 467.37 | 500.59 | 500.80 |
| Kidney | 120.74 | 93.91 | 82.64 | 153.01 | 159.63 | 158.23 |
| Liver | 191.43 | 90.36 | 84.06 | 271.66 | 285.82 | 292.68 |
| Lung | 188.86 | 97.83 | 79.77 | 199.12 | 226.95 | 207.36 |
| Lymph node | 257.20 | 68.82 | 78.16 | 486.36 | 397.09 | 495.19 |
| Mel | 102.05 | 67.47 | 61.26 | 89.21 | 88.84 | 85.16 |
| NSCLC | 107.51 | 78.64 | 73.08 | 95.62 | 92.70 | 97.06 |
| OvC | 98.24 | 53.13 | 54.49 | 68.23 | 68.35 | 67.31 |
| Ovary | 178.76 | 93.12 | 81.93 | 191.55 | 224.82 | 210.95 |
| Prostate | 110.00 | 59.28 | 61.05 | 108.29 | 100.63 | 100.68 |
| Skin | 140.62 | 89.73 | 70.81 | 173.97 | 166.38 | 166.79 |
| Spleen | 57.24 | 57.76 | 57.32 | 47.16 | 48.00 | 46.04 |
| Thyroid | 57.24 | 44.74 | 43.52 | 41.50 | 39.69 | 43.64 |
| Tonsil | 53.22 | 47.01 | 45.48 | 33.57 | 34.30 | 33.55 |

**Supplementary Table 8:** Pearson correlation results of tumor TMA and TCGA database across all panels.

| Platform (panel) Cancer Type\ | Xenium multi-tissue | Xenium breast | Xenium lung | MERSCOPE breast | MERSCOPE lung | CosMx 1k |
|---|---|---|---|---|---|---|
| Bladder cancer | 0.64 | 0.72 | 0.69 | 0.66 | NaN | 0.64 |
| Breast cancer | 0.65 | 0.66 | 0.70 | 0.71 | 0.76 | 0.66 |
| CRC | 0.75 | 0.77 | 0.76 | 0.74 | 0.73 | 0.66 |
| HNSCC | 0.76 | 0.78 | 0.77 | 0.73 | 0.68 | 0.69 |
| Melanoma | 0.65 | 0.58 | 0.63 | 0.61 | 0.59 | 0.60 |
| NSCLC | 0.69 | 0.75 | 0.59 | 0.73 | 0.52 | 0.63 |
| Ovarian cancer | 0.69 | 0.71 | 0.73 | 0.74 | 0.75 | 0.71 |

**Supplementary Table 9:** Pearson correlation results of normal TMA and GTEx database across all panels.

| Platform (panel) Tissue Type\ | Xenium multi-tissue | Xenium breast | Xenium lung | MERSCOPE breast | MERSCOPE lung | CosMx 1k |
|---|---|---|---|---|---|---|
| Bladder | 0.64 | 0.64 | 0.59 | 0.44 | 0.49 | 0.58 |
| Breast | 0.68 | 0.71 | 0.58 | 0.36 | 0.50 | 0.69 |
| Colon | 0.57 | 0.61 | 0.55 | 0.40 | 0.40 | 0.61 |
| Heart | 0.75 | 0.77 | 0.76 | 0.12 | -0.05 | 0.55 |
| Kidney | 0.72 | 0.76 | 0.74 | 0.63 | 0.69 | 0.50 |
| Liver | 0.82 | 0.85 | 0.81 | 0.56 | 0.60 | 0.69 |
| Lung | 0.74 | 0.79 | 0.69 | 0.06 | 0.18 | 0.53 |
| Lymph node | 0.41 | 0.41 | 0.32 | 0.18 | 0.38 | 0.38 |
| Ovary | 0.54 | 0.56 | 0.48 | 0.49 | 0.60 | 0.69 |
| Prostate | 0.80 | 0.81 | 0.80 | NaN | 0.72 | 0.77 |
| Skin | 0.72 | 0.82 | 0.79 | 0.37 | 0.54 | 0.68 |
| Spleen | 0.76 | 0.81 | 0.74 | 0.04 | 0.13 | 0.35 |
| Thyroid | 0.17 | 0.23 | 0.19 | 0.31 | 0.34 | 0.44 |
| Tonsil | 0.77 | 0.79 | 0.79 | 0.61 | 0.58 | 0.72 |
| Pancreas | 0.23 | 0.34 | 0.16 | NaN | 0.27 | 0.41 |

**Supplementary Table 10:** Example of gene level data.

| Core | Gene | Tissue type | Count | Code type |
|---|---|---|---|---|
| 1 | AATK | Bladder | 17 | gene |
| 1 | ABL1 | Bladder | 30 | gene |
| 1 | ABL2 | Bladder | 22 | gene |
| 1 | ACACB | Bladder | 27 | gene |
| 1 | ACE | Bladder | 21 | gene |
| 1 | ACKR1 | Bladder | 12 | gene |
| 1 | ACKR3 | Bladder | 35 | gene |
| 1 | ACKR4 | Bladder | 30 | gene |
| 1 | ACP5 | Bladder | 76 | gene |
| 1 | ACTA2 | Bladder | 31 | gene |
| 1 | ACTG2 | Bladder | 11 | gene |

**Supplementary Table 11:** Example of cell level data.

| cell_id | transcript_ counts | control_ probe_ counts | control_ codeword_ counts | unassigned_ codeword_ counts | total_counts | cell_area | nucleus_ area | x | y | core | tissue_type | geometry |
|---|---|---|---|---|---|---|---|---|---|---|---|---|
| aaaabane-1 | 213 | 0 | 0 | 0 | 213 | 116.01 | 39.78 | 7,428.87 | 21.81 | 61 | HNSCC | POINT (7428.874658203124 21.810035228729248) |
| aaaagnbb-1 | 139 | 0 | 0 | 0 | 139 | 118.67 | 10.25 | 7,428.12 | 4.89 | 61 | HNSCC | POINT (7428.116796875 4.891704702377319) |
| aaabnncg-1 | 251 | 0 | 0 | 1 | 252 | 136.64 | 71.03 | 7,437.15 | 19.53 | 61 | HNSCC | POINT (7437.15302734375 19.53349733352661) |
| aaabnnhb-1 | 107 | 0 | 0 | 0 | 107 | 60.64 | 12.51 | 7,436.43 | 10.70 | 61 | HNSCC | POINT (7436.42919921875 10.702136468887328) |
| aaabojgn-1 | 163 | 0 | 0 | 0 | 163 | 156.11 | 33.82 | 7,443.17 | 4.72 | 61 | HNSCC | POINT (7443.172753906249 4.719903254508972) |
| aaacafik-1 | 633 | 0 | 0 | 0 | 633 | 342.15 | 153.03 | 7,452.35 | 17.53 | 61 | HNSCC | POINT (7452.350927734375 17.53206386566162) |
| aaaccjoo-1 | 184 | 0 | 0 | 0 | 184 | 86.20 | 24.66 | 7,454.67 | 26.94 | 61 | HNSCC | POINT (7454.669335937499 26.943363189697266) |
| aaacdggg-1 | 89 | 0 | 0 | 0 | 89 | 60.87 | 12.24 | 7,460.32 | 29.56 | 61 | HNSCC | POINT (7460.318017578124 29.56240882873535) |
| aaacdmce-1 | 290 | 0 | 0 | 0 | 290 | 245.24 | 56.54 | 7,464.75 | 4.88 | 61 | HNSCC | POINT (7464.747314453125 4.880907201766967) |
| aaaceomi-1 | 140 | 0 | 0 | 0 | 140 | 99.52 | 28.18 | 7,466.66 | 23.29 | 61 | HNSCC | POINT (7466.6623046875 23.29304599761963) |

**Supplementary Table 12:** Example of cell by gene data.

| cell_id | core | tissue_type | ABCC11 | ADAM9 | ADGRE5 | ADH1B | … | ZEB1 | ZEB2 | ZNF562 |
|---|---|---|---|---|---|---|---|---|---|---|
| 3003241900002100003_region_1 | 124 | NSCLC | 0 | 0 | 0 | 0 | … | 0 | 0 | 0 |
| 3003241900002100004_region_1 | 124 | NSCLC | 0 | 0 | 0 | 0 | … | 0 | 0 | 0 |
| 3003241900002100006_region_1 | 124 | NSCLC | 0 | 0 | 0 | 0 | … | 0 | 0 | 0 |
| 3003241900002100007_region_0 | 61 | HNSCC | 0 | 0 | 1 | 0 | … | 0 | 0 | 0 |
| 3003241900002100007_region_1 | 124 | NSCLC | 0 | 0 | 0 | 0 | … | 0 | 0 | 0 |
| 3003241900002100008_region_1 | 124 | NSCLC | 0 | 0 | 0 | 0 | … | 0 | 0 | 0 |
| 3003241900002100009_region_0 | 61 | HNSCC | 0 | 0 | 0 | 0 | … | 0 | 0 | 0 |
| 3003241900002100010_region_0 | 61 | HNSCC | 0 | 0 | 0 | 0 | … | 1 | 2 | 0 |
| 3003241900002100010_region_1 | 124 | NSCLC | 0 | 0 | 0 | 0 | … | 0 | 0 | 0 |
| 3003241900002100011_region_0 | 61 | HNSCC | 0 | 0 | 0 | 0 | … | 0 | 0 | 0 |
